# Misoprostol regulates Bnip3 repression and alternative splicing to control cellular calcium homeostasis during hypoxic stress

**DOI:** 10.1101/313163

**Authors:** Jared T. Field, Matthew D. Martens, Wajihah Mughal, Yan Hai, Donald Chapman, Grant M. Hatch, Tammy L. Ivanco, William Diehl-Jones, Joseph W. Gordon

## Abstract

The cellular response to hypoxia involves the activation of a conserved pathway for gene expression regulated by the transcription factor complex called hypoxia-inducible factor (HIF). This pathway has been implicated in both the adaptive response to hypoxia and in several hypoxic-ischemic related pathologies. Perinatal hypoxic injury, often associated with prematurity, leads to multi-organ dysfunction resulting in significant morbidity and mortality. Using a rodent model of neonatal hypoxia and several representative cell lines, we observed HIF1α activation and down-stream induction of the cell death gene Bnip3 in brain, large intestine, and heart which was mitigated by administration of the prostaglandin E1 analog misoprostol. Mechanistically, we determined that misoprostol inhibits full-length Bnip3 (Bnip3-FL) expression through PKA-mediated NF-κB (P65) nuclear retention, and the induction of pro-survival splice variants. We observed that the dominant small pro-survival variant of Bnip3 in mouse cells lacks the third exon (Bnip3ΔExon3), whereas human cells produce a pro-survival BNIP3 variant lacking exon 2 (BNIP3ΔExon2). In addition, these small Bnip3 splice variants prevent mitochondrial dysfunction, permeability transition, and necrosis triggered by Bnip3-FL by blocking calcium transfer from the sarco/endoplasmic reticulum to the mitochondria. Furthermore, misoprostol and Bnip3ΔExon3 promote nuclear calcium accumulation, resulting in HDAC5 nuclear export, NFAT activation, and adaptive changes in cell morphology and gene expression. Collectively, our data suggests that misoprostol can mitigate the potential damaging effects of hypoxia on multiple cell types by activating adaptive cell survival pathways through Bnip3 repression and alternative splicing.

## Introduction

Hypoxia is a central element in many diseases of prematurity, including hypoxic/ischemic encephalopathy (HIE)^1^, necrotizing enterocolitis (NEC)^2^, retinopathy of prematurity^3^, and persistent pulmonary hypertension of the newborn (PPHN)^4^. In addition, cardiac dysfunction is an important predictor of morbidity and mortality in hypoxia- and asphyxia-related neonatal disorders, as impaired cardiac metabolism and contractile performance compromises tissue perfusion^5,6^. Regardless of the cause, oxygen-deprived cells display accumulating levels of transcription factors belonging to the hypoxia-inducible factor-alpha (HIFα) family. During normoxia, HIFα is hydroxylated within its oxygen degradation domain (ODD) by the prolyl-hydroxylase domain (PHD) enzymes, triggering HIFα degradation by the proteasome^7^. However, a reduced cellular oxygen tension inhibits the activity of the PHD enzymes, allowing HIFα to accumulate in the nucleus and activate transcription through dimerization with the HIFβ subunit^7^. Although cell-type specific differences in this pathway exist, there is remarkable conservation amongst multiple cell-types in response to HIFα activation, including the resulting induction in glycolytic metabolism and the reduction of mitochondrial respiration^7,8^.

HIF1α has been shown to increase the expression of members of the Bcl-2 gene family, including the BCL-2/adenovirus E1B 19 kd-interacting protein 3 (Bnip3), whose protein product plays a pivotal role in hypoxia-induced apoptosis, necrosis and autophagy^9,10^. Depending on the cellular context, Bnip3 has been previously shown to induce macro-autophagy by disrupting the Beclin-1/Bcl-2 complex^11^, promote mitochondrial outer membrane permeability (MOMP) leading to apoptosis^12,13^, and trigger mitochondrial permeability transition-dependent necrosis by releasing calcium from the endoplasmic reticulum^12,14^. In cardiomyocytes, Bnip3 expression is negatively regulated by a p65/p50 dimer of the NF-κB family (reviewed by Gordon et al^15^). Although canonical NF-κB signaling occurs through repression of Inhibitor of κB (IκB) by the IκB kinase (IKK), other signaling pathways have been shown to alter NF-κB transcriptional activity, co-factor interaction, and alter the nuclear-to-cytoplasmic shuttling of the p65 subunit^16,17^. For example, PKA phosphorylates human P65 at Serine-276 to promote nuclear accumulation and the interaction with the histone acetyl transferase p300^18–20^. However, in the context of the Bnip3 promoter, p65 serves to recruit HDAC1 to repress gene expression^15^.

Bnip3 has been shown to be alternatively spliced leading to the production of an endogenous inhibitor that lacks the third exon, called Bnip3ΔExon3^21^. The fusion of exon 2 to exon 4 of the *Bnip3* gene results in a frame-shift, a premature stop codon, and the production of a truncated protein with a divergent C-terminus. Bnip3ΔExon3 appears to act as an endogenous inhibitor of full-length Bnip3 (Bnip3-FL) by preventing mitochondrial depolarization, and promoting cell viability^21^. However, the precise mechanism(s) by which Bnip3ΔExon3 inhibits hypoxia- and Bnip3-induced cell death remain less clear.

Recently, we demonstrated that Bnip3 expression was elevated in enterocytes subjected to nutrient/oxidative stress induced by breast milk fortifiers, while Bnip3-induced enterocyte cell death was inhibited by exogenous expression of Bnip3ΔExon3^22^. Furthermore, fortifier-induced cellular toxicity was completely abrogated by treatment of enterocytes with the prostaglandin analog misoprostol^22^. These compelling findings led us to investigate whether misoprostol could protect cells against hypoxia-induced injury. Furthermore, given the degree of conservation in the cellular response to hypoxia, we sought to determine if misoprostol could protect multiple cell types from Bnip3-induced injury, such as that occurring during neonatal hypoxia/asphyxia.

In this report, we provide evidence that misoprostol opposes hypoxia-induced Bnip3 expression in multiple tissues, including gut, brain, and the heart. In cultured cells, we observed that misoprostol activates PKA signaling and promotes nuclear localization of P65 to suppress Bnip3-FL expression and increase the expression of smaller splice variants. In addition, we discovered a previously unidentified Bnip3 splice variant lacking exon 2 (BNIP3ΔExon2), which is expressed in human cells. Remarkably, this spice variant contains the same frame-shift as mouse Bnip3ΔExon3, resulting a conserved C-terminal amino acid sequence. Mechanistically, we determined that a combination of NF-κB and HIF1α activation alters BNIP3 splicing and induces the expression of the smaller variants. Finally, we experimentally altered the ratio of Bnip3-FL to Bnip3ΔExon3, and Bnip3-FL to BNIP3ΔExon2 in multiple cell types, and observed that the smaller variants prevent mitochondrial permeability transition and cell death by inhibiting mitochondrial calcium accumulation and promoting nuclear calcium-dependent cell adaptation. These findings suggest that prostaglandin signaling may play an important developmental role that can be used therapeutically to circumvent many of the morbidities associated with neonatal hypoxia.

## Results

### Misoprostol inhibits Bnip3-FL expression in vivo

Previously, we demonstrated that misoprostol could inhibit cell death in cultured enterocytes exposed to nutrient stress^22^. Moreover, our previous work implicated the hypoxia-inducible gene Bnip3 in this process. In order to further these studies *in vivo*, we exposed neonatal rats to 7 days of hypoxia (10% oxygen), prior to weaning, with and without misoprostol treatment (10 μg/kg; subcutaneous). This protocol has been previously shown to induce cognitive impairment consistent with HIE^23,24^. Bnip3 expression analysis by western blot demonstrated increased Bnip3-FL expression in the hippocampus, large intestine, and heart of hypoxia-exposed animals, which was completely abrogated by concurrent misoprostol treatment (Figure 1A).

**Figure 1.**
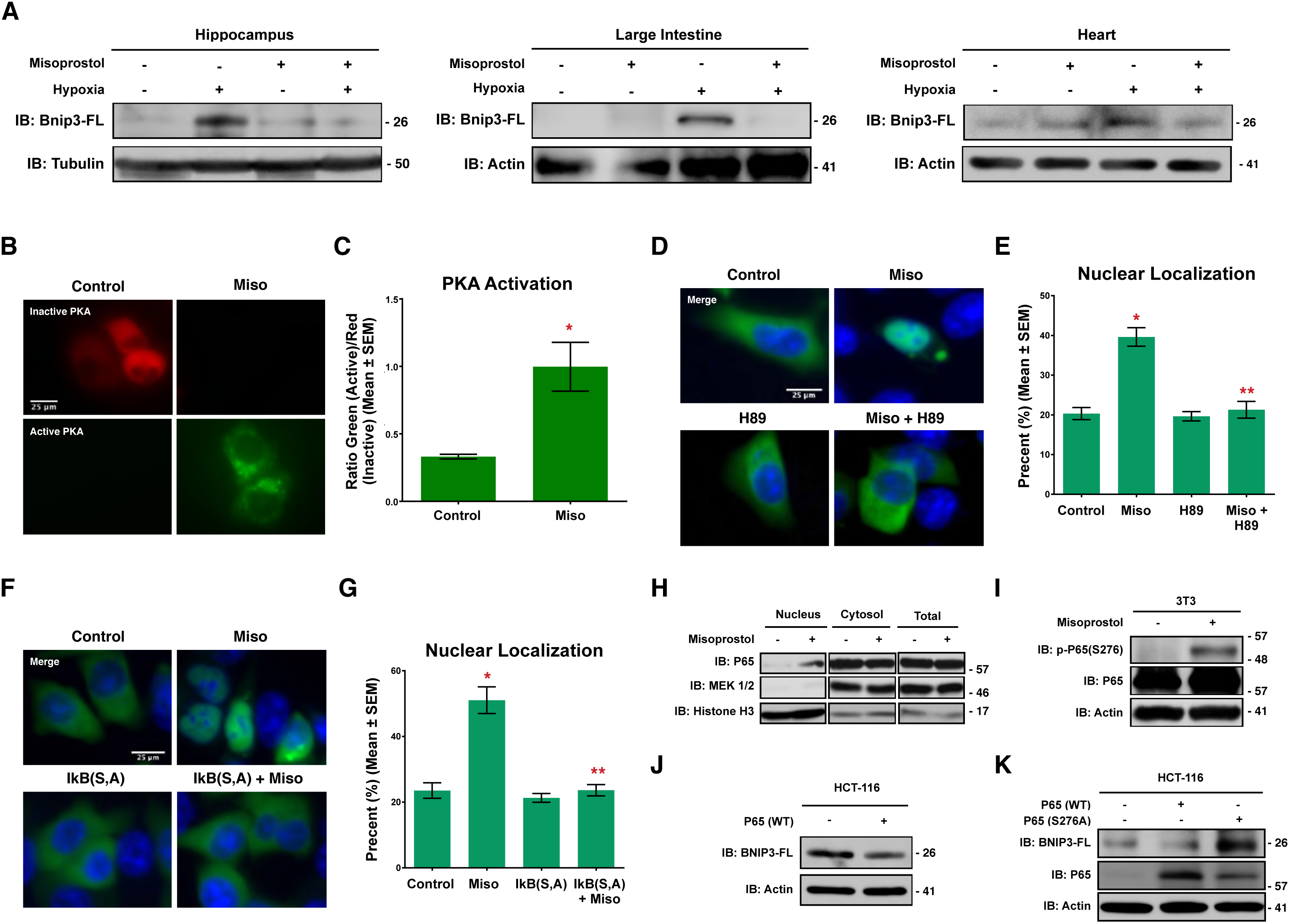
Misoprostol opposes hypoxia-induced Bnip3-FL expression. (**A**) Immunoblot for Bnip3-FL in protein extracts from the hippocampus, large intestine and heart of PND10 rat pups exposed to hypoxia (10% O_2_) ± misoprostol for 7 days. (**B**) HCT-116 cells were transfected with PKA biosensor (pPHT-PKA), and treated with 10 μM misoprostol or vehicle for 2hrs. Cells were imaged by standard fluorescence microscopy. (**C**) Quantification of fluorescent images in (B) by measuring the ratio of green (active) to red (inactive) fluorescent signal, normalized to cell area, quantified in 10 random fields. (**D**) HCT-116 cells were transfected with GFP-P65 (Green) and were treated with 10 μM misoprostol ± 10 μM H89 for 20hrs. Cells were stained with Hoechst (blue) and imaged by standard fluorescence microscopy. (**E**) Quantification of fluorescent images in (D) by calculating the percentage of cells with nuclear P65 signal over 10 random fields. (**F**) HCT-116 cells were transfected with IkBa(SS32,36AA) [indicated as IkB (S, A)] or an empty vector control. GFP-P65 (Green) was used to indicate subcellular P65 localization in all conditions. Cells were treated with 10 μM misoprostol or vehicle control for 20hrs. Cells were stained with Hoechst (blue) and imaged by standard fluorescence microscopy. (**G**) Quantification of fluorescent images in (D) by calculating the percentage of cells with nuclear P65 signal over 10 random fields. (**H**) HCT-116 cells were treated with 10 μM misoprostol or vehicle for 20hrs, during extraction proteins were fractioned according to their sub-cellular compartment. Protein extracts were immunoblotted, as indicated. (**I**) 3T3 cells were treated with 10 μM misoprostol or vehicle control for 4 hours. Protein extracts were immunoblotted as indicated. (**J**) HCT-116 cells were transfected with wild-type P65 or empty vector control. Extracts were immunoblotted, as indicated. (**K**) HCT-116 cells were transfected with P65 constructs for 20hr. Extracts were immunoblotted, as indicated. Data are represented as mean ± S.E.M. *^*^P*<0.05 compared with control, while *^**^P*<0.05 compared with treatment, determined by 1-way ANOVA or by unpaired t-test.

Canonical NF-κB signaling has been previously shown to be a critical regulator of Bnip3 expression during cardiac hypoxia (reviewed by Gordon et al^15^). Moreover, protein kinase-A (PKA) has been shown to phosphorylate P65 of the NF-κB complex at Serine-276 to promote nuclear accumulation^18^. Thus, we tested the hypothesis that misoprostol could repress Bnip3-FL expression through a conserved PKA- and NF-κB-dependent mechanism. Using a genetically-encoded plasmid-based PKA biosensor (pPHT-PKA)^25^, we observed marked PKA activation in HCT-116 cells treated with misoprostol (Figure 1B, C). Consistent with our hypothesis, we observed increased nuclear localization of a human P65-GFP construct in misoprostol treated HCT-116 cells (Figure 1D, E). However, when cells were co-treated with misoprostol and the PKA-inhibitor H89, this effect was lost and P65 was retained in the cytoplasm. To further investigate the role of NF-κB signalling in this misoprostol-induced response, we expressed the same P65-GFP construct concurrently with a dominant negative form of the Inhibitor of kappa-B (IκB S32A, S36A or IκB-SA), in misoprostol treated HCT-116 cells. Here we observed that the misoprostol-induced nuclear localization was lost in the presence of IκB-SA (Figure 1F, G). Additionally, we performed nuclear and cytosolic fractionation studies to biochemically confirm the observed phenomenon of P65-nuclear localization. In HCT-116 cells treated with a vehicle control or misoprostol, we observed that misoprostol treatment had no effect on total P65 expression (Figure 1H). However, consistent with our P65-GFP findings, we observed an increase in levels of nuclear P65. Next, we assessed the role of misoprostol in the regulation of P65 phosphorylation at a known PKA phosphorylation site, serine-276. Shown in Figure 1I, we observed that misoprostol treatment increased phospho-p65. The migration of this band is consistent with information provided by the manufacturer.

To confirm that NF-κB regulates BNIP3-FL expression in HCT-116 cells, we expressed P65 in this cell line and analyzed BNIP3-FL expression by western blot. Ectopic expression of P65 reduced BNIP3-FL protein levels (Figure 1J). Furthermore, we expressed wild-type P65, and a P65 mutant form, where the PKA phospho-acceptor site, Ser-276, was mutated to a neutral alanine (P65-S276A) thereby ablating phosphorylation at this site. This experiment confirmed that wild-type P65 represses BNIP3FL expression; however, when Ser-276 is neutralized BNIP3-FL expression is enhanced (Figure 1K).

Collectively, these findings support a model of Bnip3-FL repression where misoprostol activates PKA signaling to promote P65 phosphorylation at the Ser-276 site, leading to nuclear accumulation of the NF-κB complex and subsequent repression of BNIP3-FL expression.

### Discovery of BNIP3ΔExon2

Recently, it was reported that the rat *Bnip3* pre-mRNA undergoes alternative splicing to yield a full-length protein containing all six exons, and a small splice variant lacking the third exon (Bnip3ΔExon3)^21^. Moreover, the fusion of exon 2 to exon 4 in the Bnip3ΔExon3 mRNA results in a frame-shift and the generation of a pre-mature stop codon. Functionally, Bnip3ΔExon3 acts as an endogenous inhibitor of BNIP3-FL, blocking hypoxia- and nutrient stress-induced cell death^21,22^. Based on our findings that misoprostol supressed BNIP3-FL expression elicited by neonatal hypoxia, we used semi-quantitative RT-PCR to evaluate whether misoprostol could alter *Bnip3* splicing. Using primers that spanned the exon 2-4 junction, we detected a smaller amplicon representing Bnip3ΔExon3 in RNA isolated from hearts of neonatal rats treated with misoprostol, but not in vehicle treated animals (Figure 2A).

**Figure 2.**
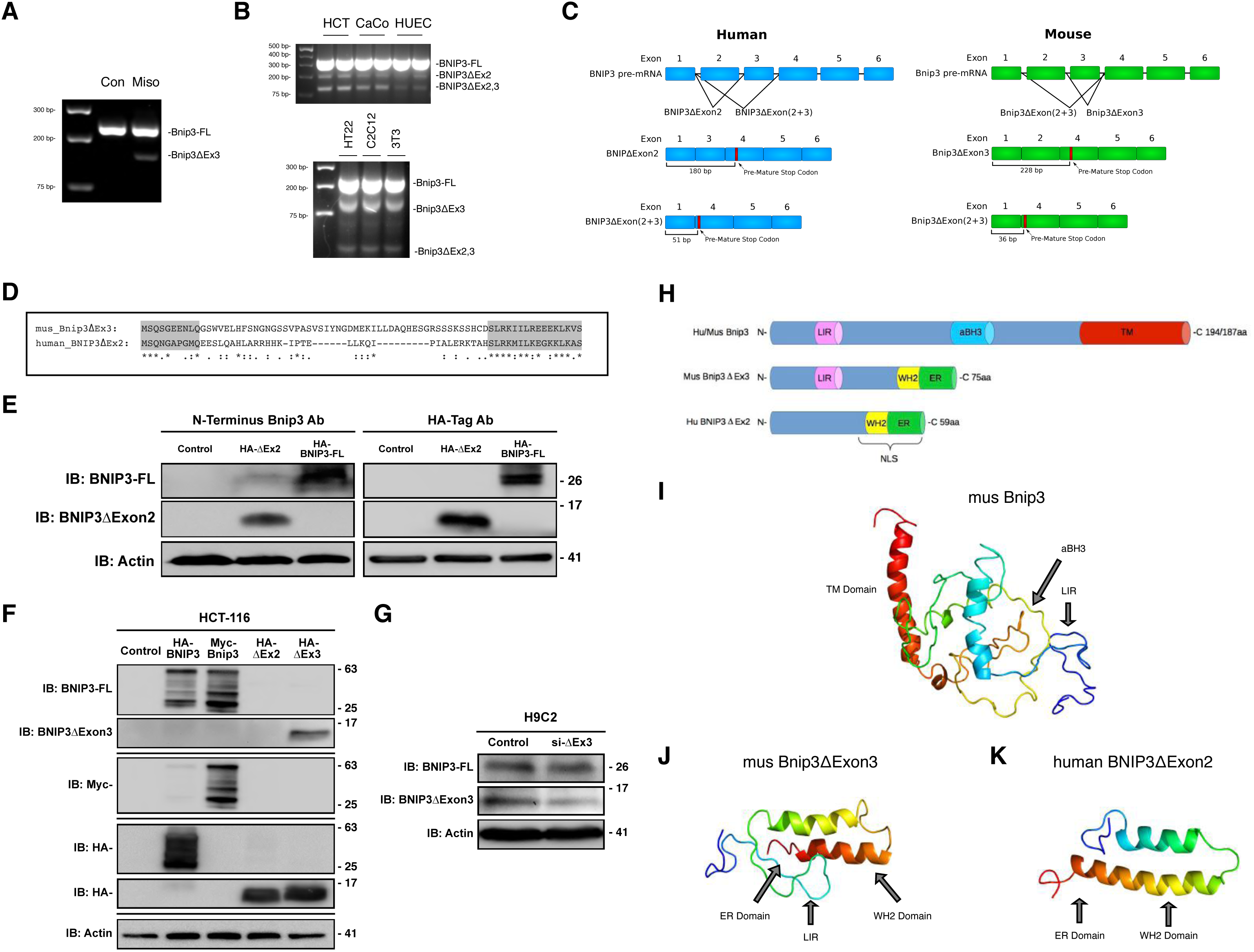
Misoprostol alters the splicing of Bnip3. (**A**) RT-PCR for Bnip3-FL and Bnip3ΔExon3 in the heart of PND10 rat pups treated with misoprostol for 7 days. (**B**) RT-PCR for Bnip3 isoforms comparing expression patterns between human and mouse cell lines. (**C**) Bnip3 splicing diagram showing BNIP3ΔExon2 and Bnip3ΔExon3, indicating the differences in splicing between human and mouse. (**D**) Amino acid sequence alignment for human BNIP3ΔExon2 and mouse Bnip3ΔExon3. (**E**) Immunoblot showing that a commercially available antibody targeted to the amino-terminus (N-Terminus) of BNIP3 (CST #44060) can detect overexpressed HA-BNIP3-FL and HA-BNIP3ΔExon2. (**F**) Immunoblot demonstrating the specificity of our custom antibody targeted to the amino-terminus (N-Terminus) of BNIP3, where the antibody is able to detect overexpressed HA-BNIP3-FL (Human), Myc-BNIP3-FL (Mouse) and HA-BNIP3ΔExon3 (Mouse), but not HA-BNIP3ΔExon2 (Human). (**G**) Immunoblot demonstrating the specificity of si-BNIP3ΔExon3 (indicated as si-ΔEx3), shown using our custom antibody targeted to the amino-terminus (N-Terminus) of BNIP3. (**H**) Linear motif diagram for Bnip3-FL, human BNIP3ΔExon2, and mouse Bnip3ΔExon3. (**I**) Model of *Mus* Bnip3-FL indicating locations of the transmembrane (TM) domain, atypical BH3 domain (aBH3) and LC3-Interacting region (LIR) present on this isoform. (**J**-**K**) Comparison of *Mus* Bnip3ΔExon3 and human BNIP3ΔExon2, indicating the locations of the ER domains, WH2 domains and the *Mus* Bnip3ΔExon3 specific LC3-Interacting region (LIR).

To our knowledge, Bnip3ΔExon3 expression has not been reported at the protein level. Recently, a commercially available antibody targeting exon 1 of BNIP3 has come to market. This antibody could hypothetically detect smaller BNIP3 variants; however, this antibody only detects human BNIP3 protein (Cell Signaling Technology; #44060). In order to evaluate the role of misoprostol promoting BNIP3 splicing in human cell lines, we attempted to clone human BNIP3ΔExon3. We designed a forward primer encompassing the 5’-ATG of human *BNIP3* and a reverse primer containing the premature stop codon identified in mouse Bnip3ΔExon3. Restriction enzyme sites were incorporated into these primers to allow easy ligation into pcDNA3 and sequencing. Interestingly, in three distinct human cells lines we observed three amplicons generated by these primers (Figure 2B). Based on sequencing analysis, the largest amplicon represents a fragment of BNIP3-FL. However, the middle and smallest amplicons represent splice variants lacking exon 2 (BNIP3ΔExon2) and exons 2 and 3 (BNIP3ΔExon2+3), respectively. Using the same approach in three representative mouse cell lines, we identified a fragment of BNIP3-FL, Bnip3ΔExon3, and Bnip3ΔExon2+3 (Figure 2B). These findings suggest that Bnip3 is differentially spliced in human versus rodent cells (Figure 2C). The variants with two missing exons form very small peptides due to early stop codons in their reading frame (Figure 2C and Supplemental). However, the mouse Bnip3ΔExon3 and human BNIP3ΔExon2 share the same frame shift, and generate a conserved C-terminus when translated (Figure 2D) (Alignment performed on the CLUSTAL O 1.2.4 multiple sequence alignment tool). Importantly, when expressed in HCT-116 cells, we were able to detect both human HA-BNIP3-FL and HA-BNIP3ΔExon2 with the commercially available N-terminal BNIP3 antibody (Figure 2E), suggesting that detection of endogenous protein of these small BNIP3 splice variants is possible in human cells and tissue. However, this commercially-available antibody could not detect mouse Bnip3ΔExon3 protein. To solve this, we designed a custom Bnip3 antibody specifically targeted to a region of the N-terminus of full-length Bnip3 that is conserved in humans and rodents. We then performed a series of validation experiments to test this custom antibody. First, we tested if the antibody was able to detect HA- or myc-tagged versions of BNIP3-FL and Bnip3∆Exon3 in a side-by-side comparison with commercial HA-tag or myc-tag antibodies. As shown in Figure 2F, the antibody successfully detects both human and mouse full length BNIP3, and what is likely the monomeric and dimeric forms of BNIP3-FL at the approximate sizes of 25 and 63 kDa, respectively. Importantly, we could detect the smaller mouse isoform, Bnip3ΔExon3, but not the human isoform, BNIP3ΔExon2, as this epitope spans the second exon in the human gene. Second, we used an siRNA targeted to the unique sequence of Bnip3∆Exon3 to knockdown endogenous Bnip3ΔExon3 protein (Figure 2G). This set of experiments provides evidence that both the custom antibody successfully detects Bnip3ΔExon3, and that the siRNA knockdown of Bnip3ΔExon3 shows specificity for Bnip3ΔExon3 and has little effect on the expression of full-length Bnip3 protein. To our knowledge this is the first documentation of Bnip3ΔExon3 detected as a protein; furthermore, we demonstrated endogenous expression of this protein in a rodent cardiac cell line.

*In silico* domain mapping of the smaller BNIP3 variants revealed that neither mouse Bnip3ΔExon3 nor human BNIP3ΔExon2 contain the atypical BH3 domain of the full-length splice variants, and only mouse Bnip3ΔExon3 retained the N-terminal LC3 interacting region (LIR) (Figure 2H). Interestingly, both mouse Bnip3ΔExon3 and human BNIP3ΔExon2 contain C-terminal WH2 actin binding motifs and a di-lysine ER retention signal (ER). Moreover, the C-terminal region of human BNIP3ΔExon2 contains a predicted bipartite nuclear localizing signal (NLS). Finally, the Phyre2 (**P**rotein **H**omology/analog**Y R**ecognition **E**ngine V 2.0) web portal for protein modeling provided the predicted structure of the smaller Bnip3 splice variants, described previously^26,27^. We used the predicted structure of mouse Bnip3-FL for comparison (Figure 2I). Intriguingly, the Phyre2 portal predicted a remarkably similar structure for both mouse Bnip3ΔExon3 and human BNIP3ΔExon2 (Figure 2J, K), which supports the idea that they may function in a similar biological manner.

### HIF1α and NF-κB P65 regulate BNIP3 expression and splicing

To further investigate how the expression of the smaller BNIP3 variants are regulated, we treated HCT-116 with misoprostol, and the HIFα stabilizer cobalt chloride (CoCl_2_) for 16-hours. Using semi-quantitative RT-PCR and primers designed to detect both BNIP3-FL and BNIP3ΔExon2, we observed that misoprostol alone reduced BNIP3-FL expression, while CoCl_2_ treatment induced the expression of both BNIP3 splice variants (Figure 3A). However, in the presence of misoprostol, we observed proportionally more BNIP3ΔExon2. Next, we preformed the same experiment using the N-terminal BNIP3 antibody, combined with western blot analysis, and observed induction of BNIP3-FL concurrent with increased HIF1α when treated with CoCl_2_ (Figure 3B). When combined with misoprostol, CoCl_2_ induced the expression of a 11-17 kDa band as detected by the N-terminal BNIP3 antibody, consistent with migration pattern of BNIP3ΔExon2 observed in Figure 2E. Interestingly, at this dose and time point misoprostol did not affect BNIP3-FL protein expression. To reconcile this observation with our misoprostol findings *in vivo*, we extended our time-course and treated HCT-116 cells with CoCl_2_ and misoprostol for 36-hours. At this time point misoprostol attenuated CoCl_2_ induced BNIP3-FL expression, while the expression BNIP3ΔExon2 was less effected (Figure 3C). These findings suggest that misoprostol alters BNIP3 splicing at earlier time points then represses BNIP3-FL expression at later time points. Next, we performed a gain-of-function experiment in HCT-116 cells to assess the expression of BNIP3 splice variants by western blot. Shown in Figure 3D, expression of HIF1α induced the expression of BNIP3-FL; however, when HIF1α expression was combined with NF-κB P65, BNIP3ΔExon2 expression was increased and BNIP3-FL expression was proportionally decreased.

**Figure 3.**
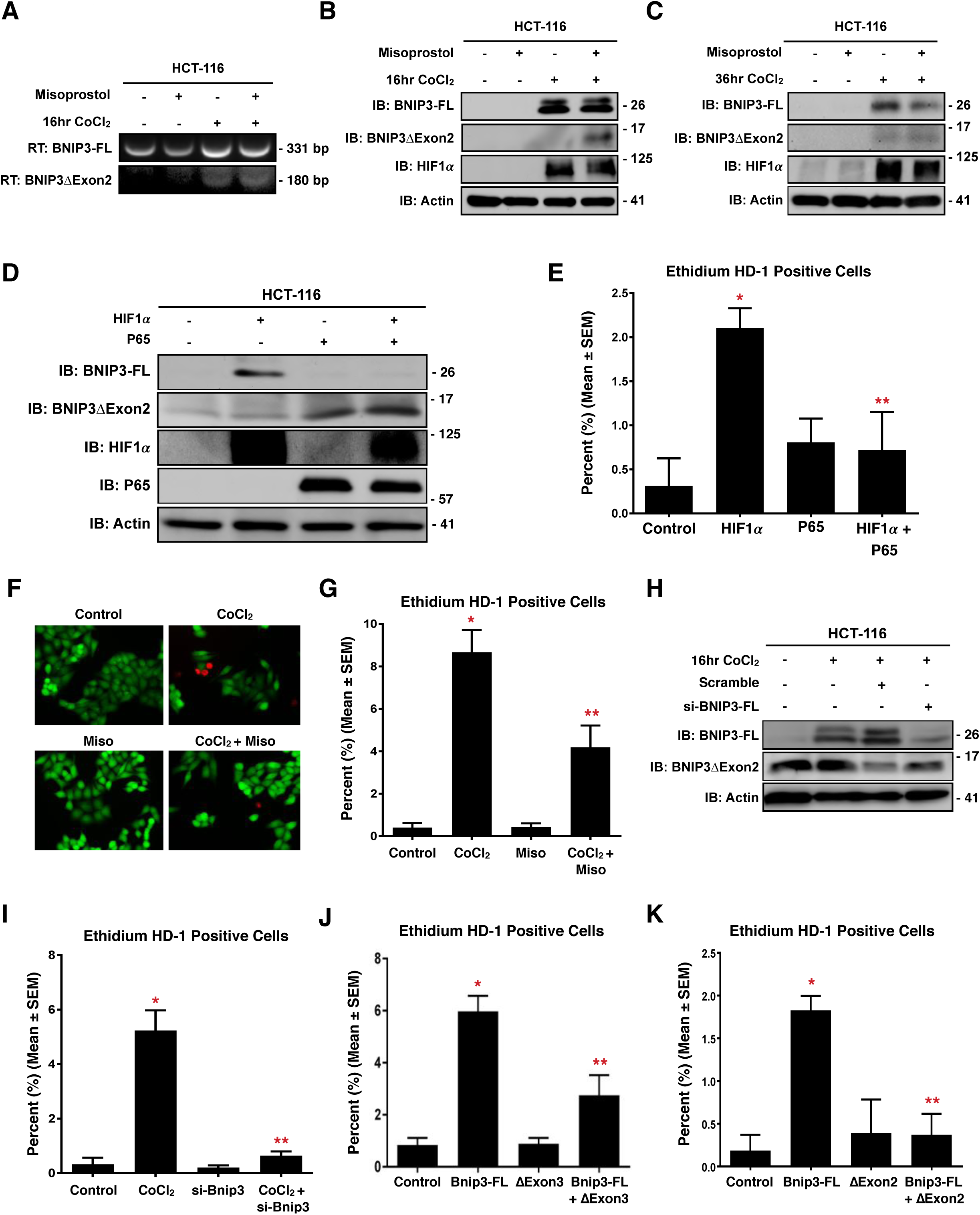
HIF1α and P65 drive expression of pro-survival BNIP3 splice variants. (**A**) HCT-116 cells were treated with 200 μM cobalt chloride ± 10 μM misoprostol or vehicle control for 20hrs. RNA was isolated and RT-PCR was performed for BNIP3 isoforms. (**B**) HCT-116 cells were treated as in (A). Protein extracts were immunoblotted, as indicated. (**C**) Immunoblot of protein extracts from HCT-116 cells treated with misoprostol and CoCl_2_ in for 36hrs. (**D**) HCT-116 cells were transfected with HIF1α ± P65. Protein extracts were immunoblotted, as indicated. (**E**) Quantification calcein-AM and ethidium homodimer-1 stained HCT-116 cells transfected with HIF1α and/or P65. (**F**) HCT-116 cells treated with 200 μM cobalt chloride ± 10 μM misoprostol or vehicle control for 20hrs. Live cells were stained were stained with calcein-AM (green)and necrotic cells were stained with ethidium homodimer-1 (red), cells were imaged by standard fluorescence microscopy. (**G**) Fluorescent images were quantified by calculating the percent of necrotic cells (ethidium homodimer-1 positive) cells in 10 random fields. (**H**) Immunoblot of HCT-116 cells transfected with si-BNIP3-FL or scrambled control. Cells were treated with 200 μM cobalt chloride or vehicle control for 20hrs. (**I**) HCT-116 cells treated as in (H) and cells were stained as in (F), and quantified as indicated in (G). (**J**) Quantification of HCT-116 cells transfected with Bnip3-FL, Bnip3ΔExon3 or empty vector control. Cells were stained, and quantified as indicated in (F). (**K**) Quantification of HCT-116 cells transfected with Bnip3-FL, BNIP3ΔExon2 or an empty vector control. Cells were stained, and quantified as indicated in (F). Data are represented as mean ± S.E.M. *^*^P*<0.05 compared with control, while *^**^P*<0.05 compared with treatment, determined by 1-way ANOVA.

To evaluate the functional consequences of altered BNIP3 splicing, we transfected HCT-116 cells with HIF1α and NF-κB P65, and assessed cell viability by using calcein-AM, which stains viable cells green, and ethidium homodimer-1, which stains necrotic cells red. HIF1α expression increased the percentage of ethidium positive (necrotic) cells, whereas the combination of both HIF1α and NF-κB P65 had no effect on cell death (Figure 3E). Next, we treated HCT-116 cells with CoCl_2_, to induce a HIF/BNIP3 response, with or without the addition of misoprostol. Overnight treatment of cells with CoCl_2_ significantly increased the percentage of ethidium positive cells (Figure 3F, G). However, when cells were treated with both CoCl_2_ and misoprostol, the percentage of ethidium positive cells was significantly reduced. To evaluate the role of BNIP3-FL in CoCl_2_-induced cell death, we performed Bnip3-FL knockdown studies. Prior to functional assays, we transfected HCT-116 cells with si-BNIP3-FL, and treated with CoCl_2_. We observed that the siRNA reduced BNIP3-FL expression without affecting BNIP3ΔExon2 expression (Figure 3H). We then assessed the functional consequences of this knockdown, by demonstrating that that si-BNIP3-FL is sufficient to prevent CoCl_2_-induced necrotic cell death in HCT-116 cells (Figure 3I). Furthermore, in parallel gain-of-function experiments, we transfected HCT-116 cells with Bnip3-FL to induced necrosis, and added either Bnip3ΔExon3 or BNIP3ΔExon2. Shown in Figure 3J and 3K, Bnip3-FL increased the percentage of ethidium positive cells, which was reduced by co-expression of either Bnip3ΔExon3 or BNIP3ΔExon2. These findings suggest that both Bnip3ΔExon3 and BNIP3ΔExon2 act as endogenous inhibitors of Bnip3-FL-induced cell death.

### Misoprostol and Bnip3ΔExon3 protect against hypoxia-induced mitochondrial dysfunction

Clinical evidence suggests that cardiac dysfunction, and the presence of serum cardiac marker-proteins, are important predictors of morbidity and mortality during neonatal hypoxia/asphyxia^5,6^. Moreover, recent experimental evidence has implicated mitochondrial permeability transition, rapid dissipation of the mitochondrial membrane potential, and mitochondrial superoxide production as pre-cursor events leading to regulated necrosis^28,29^. Thus, we evaluated the effect of misoprostol and Bnip3ΔExon3 on hypoxia-induced mitochondrial dysfunction in primary neonatal ventricular myocytes. First, we assessed mitochondrial membrane potential by TMRM staining. Shown in Figure 4A and B, exposure to hypoxia (10% O_2_ for 48-hours) reduced mitochondrial membrane potential, which was completely attenuated by treatment with misoprostol. Next, we evaluated mitochondrial super oxide levels with the dye MitoSOX, and observed that hypoxia-induced super oxide accumulation was prevented by misoprostol treatment (Figure 4C, D). We also evaluated mitochondrial respiration in primary ventricular myocytes using a metabolic flux analyzer (Seahorse, XF24). Consistent with our TMRM observations, 48-hours of hypoxia exposure reduced both basal and maximal respiration, which was completely abrogated by misoprostol treatment (Figure 4E, F). Finally, we generated a Bnip3ΔExon3 lentivirus and evaluated if this Bnip3 splice variant could overcome mitochondrial perturbations induced by hypoxia exposure. We found that primary myocytes transduced with Bnip3ΔExon3 lentivirus gained resistance to mitochondrial depolarization caused by hypoxia (Figure 4G, H). Collectively, these findings demonstrate that both misoprostol and Bnip3ΔExon3 confer some degree of protection against hypoxia-induced mitochondrial dysfunction in primary neonatal cardiomyocytes.

**Figure 4.**
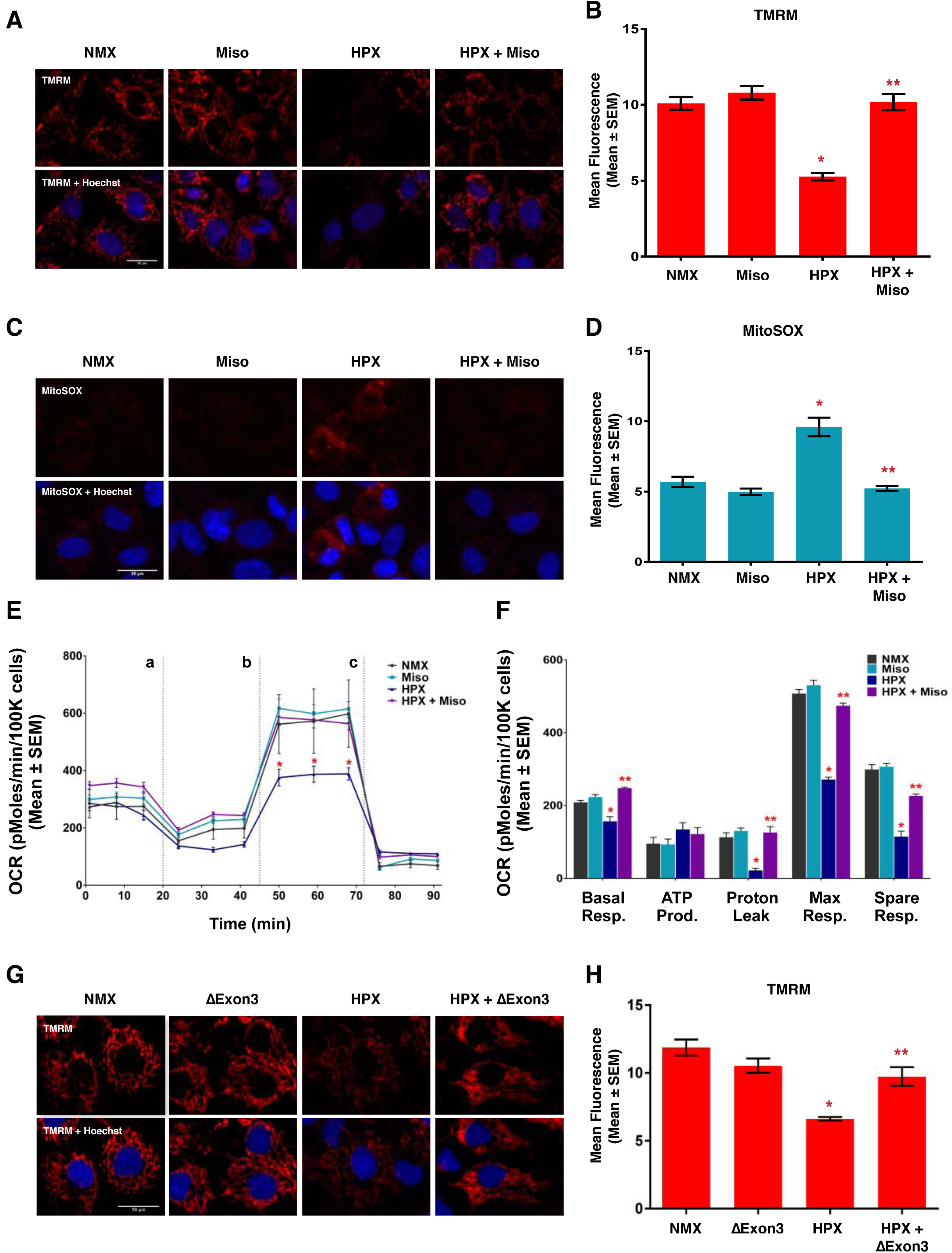
Misoprostol opposes hypoxia-induced mitochondrial dysfunction in primary ventricular neonatal cardiomyocytes. (**A**) Primary ventricular neonatal cardiomyocytes (PVNM) cells were treated with 10 μM misoprostol (Miso) ± 10% O_2_ (HPX) for 48 hours. Cells were stained with TMRM (red) and Hoechst (blue) and imaged by standard fluorescence microscopy. (**B**) Quantification of TMRM in (A), red fluorescent signal was normalized to cell area and quantified in 10 random fields. (**C**) PVNM cells were treated as described in (A). Cells were stained with MitoSOX (red) to evaluate mitochondrial superoxides and Hoechst (blue) and imaged by standard fluorescence microscopy. (**D**) Quantification of (C), red fluorescent signal was normalized to cell area and quantified in 10 random fields. (**E**) Oxygen consumption rate (OCR) was evaluated on a Seahorse XF-24 in PVNM. To evaluate mitochondrial function, wells were injected with oligomycin (1 μM) (a), FCCP (1 μM) (b), and antimycin A (1 μM) and rotenone (1 μM) (c). (**F**) Calculated respiration rates from (E). (**G**) PVNM cells were transduced with Bnip3ΔExon3 ± 10% O_2_ (HPX) for 48 hours. Cells were stained with TMRM (red) and Hoechst (blue) and imaged by standard fluorescence microscopy. (**H**) Quantification of (G), red fluorescent signal was normalized to cell area and quantified in 10 random fields. Data are represented as mean ± S.E.M. *^*^P*<0.05 compared with control, while *^**^P*<0.05 compared with hypoxia treatment, determined by 1-way ANOVA.

### Bnip3-FL-induced mitochondrial perturbations are inhibited by the small Bnip3 splice variants

Next, we evaluated how misoprostol and Bnip3-FL affect mitochondrial membrane potential in human (HCT-116) and rodent (H9c2) cell lines. Staining with TMRM, we observed that CoCl_2_ significantly reduced mitochondrial membrane potential; however, co-treatment with misoprostol completely abolished this effect (Figure 5A, B). We assessed mitochondrial membrane potential in cultured H9c2 cells, where CMV-GFP was used to identify transfected cells. Shown in Figure 5C, knockdown of Bnip3∆Exon3 reduced the effectiveness of misoprostol in moderating the depolarization caused by treatment with CoCl_2_. Therefore, we decided to directly examine the effects of the smaller splice variants on membrane potential in transfection experiments. Bnip3-FL expression reduced TMRM staining, where co-expression of Bnip3ΔExon3 and Bnip3-FL restored TMRM staining to levels comparable to control (Figure 5D), and BNIP3ΔExon2 had the same effect (Figure 5E, F). Next, we assessed mitochondrial permeability transition by staining H9c2 cells with calcein-AM, where cobalt chloride is used to quench the cytosolic emission^30^. With this technique, loss of mitochondrial puncta is interpreted as permeability transition. A CMV-dsRed plasmid was included to identify transfected cells. Expression of Bnip3-FL in H9c2 cells resulted in a significant decrease in the number of transfected cells with observable puncta; whereas, co-expression of either Bnip3ΔExon3 (Figure 5G, H) or BNIP3ΔExon2 (Figure 5I) restored the number of punctate cells. Finally, to evaluate whether these mitochondrial perturbations translated into an effect on cell viability, we transfected H9c2 cells with Bnip3-FL and with and without Bnip3ΔExon3, then assessed the percentage of ethidium homodimer positive cells. As shown in Figure 5J and K, Bnip3-FL-induced cell death was reversed in the presence of Bnip3ΔExon3 in this cardiac cell line.

**Figure 5.**
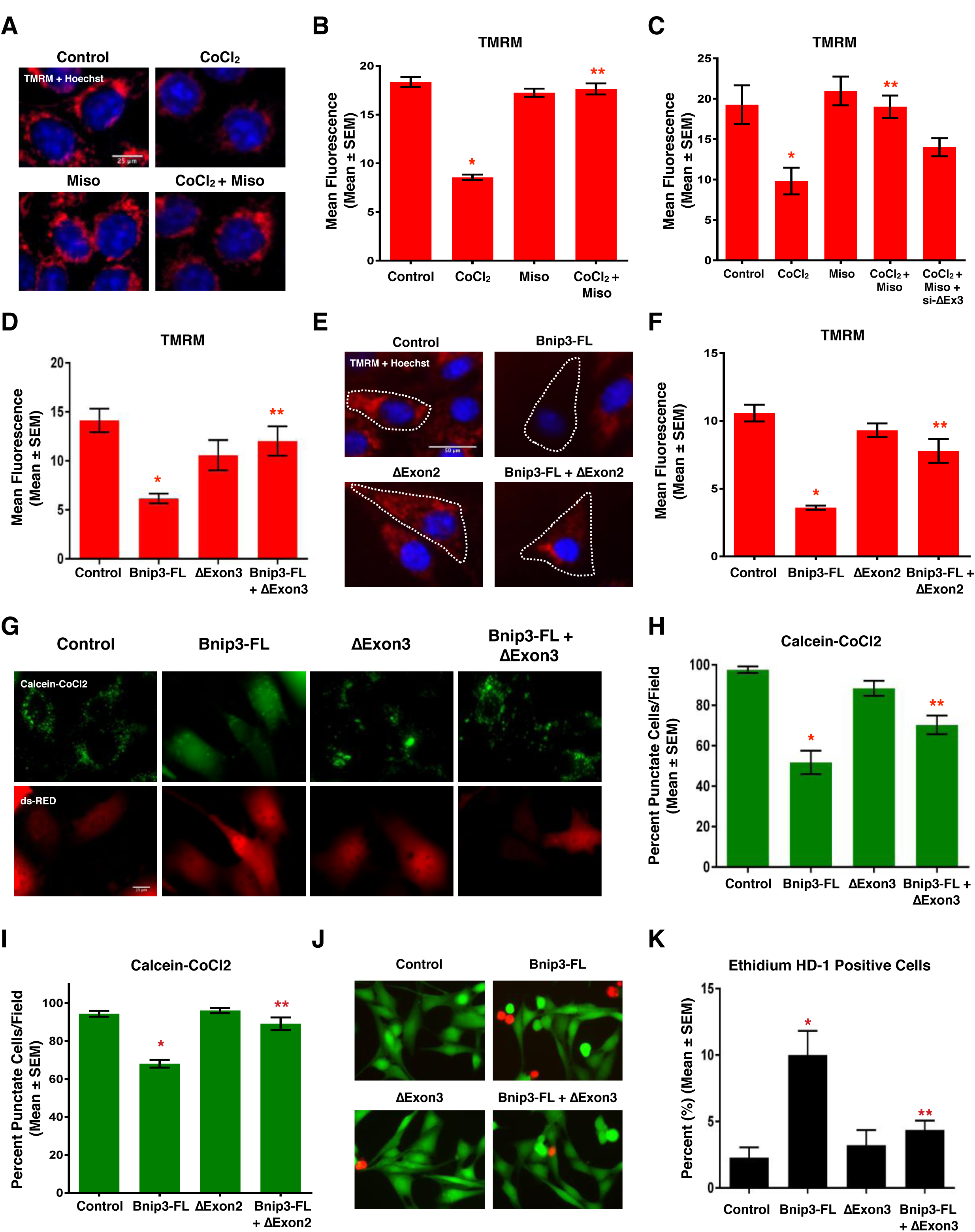
Bnip3 splice variants oppose mitochondrial perturbations. (**A**) HCT-116 cells treated with 200 μM cobalt chloride ± 10 μM misoprostol or vehicle control for 20hrs. Cells were stained with TMRM (red) and Hoechst (blue) and imaged by standard fluorescence microscopy. (**B**) Quantification of TMRM in (A), red fluorescent signal was normalized to cell area and quantified in 10 random fields. (**C**) Quantification of H9c2 cells treated with 200 μM cobalt chloride ± 10 μM misoprostol or vehicle control for 20hrs, and transfected with si-Bnip3ΔExon3 or a scrambled control. Cells were stained as in (A) and quantified as in (B). (**D**) Quantification of H9c2 cells were transfected with Bnip3-FL, Bnip3ΔExon3 or an empty vector control. TMRM staining was quantified as in (B). (**E**) H9c2 cells were transfected with Bnip3-FL, BNIP3ΔExon2 or empty vector control. Outlines indicate CMV-GFP positive cells, included to identify transfected cells. Cells were stained as in (A). (**F**) Quantification of (E), red fluorescent signal was normalized to cell area and quantified in 10 random fields. (**G**) H9c2 cells were transfected with Bnip3-FL, Bnip3ΔExon3 or empty vector control. CMV-dsRed (red) was used to identify transfected cells. Cells were stained with calcein-AM and cobalt chloride (CoCl_2_, 5 μM) to assess permeability transition. (**H**) Quantification of (G) by calculating the percentage of cells with punctate calcein signal in 10 random fields. (**I**) Quantification of H9c2 cells transfected with Bnip3-FL, BNIP3ΔExon2 or empty vector control. CMV-dsRed was used to identify transfected cells. Cells were stained and quantified as indicated in (H). (**J**) H9c2 cells transfected with Bnip3-FL, BNIP3ΔExon3 or empty vector control. Live cells were stained with calcein-AM (green), and necrotic cells were stained with ethidium homodimer-1 (red), cells were imaged by standard fluorescence microscopy. (**K**) Fluorescent images in (J) were quantified by calculating the percent of necrotic cells (ethidium homodimer-1 positive) cells in 10 random fields. Data are represented as mean ± S.E.M. *^*^P*<0.05 compared with control, while *^**^P*<0.05 compared with Bnip3-FL treatment, determined by 1-way ANOVA.

### Bnip3 splice variants regulate mitochondrial calcium homeostasis

An important component of regulated necrosis involves permeability transition triggered by elevations in mitochondrial calcium^29,31,32^, where calcium release from the endoplasmic reticulum can serve as a trigger for permeability transition^33,34^. To more fully investigate the role of cellular calcium in Bnip3-regulated permeability transition, we used organelle-targeted plasmid-based calcium biosensors, called GECOs (<u>G</u>enetically <u>E</u>ncoded <u>C</u>a^2+^ indicators for <u>O</u>ptical imaging)^35^. We used the mitochondrial matrix-targeted red GECO, known as mito-carmine (mito-Car-GECO) and ER-targeted LAR-GECO (ER-LAR-GECO)^36^ to test the hypothesis that the effects of Bnip3 splice variants on mitochondrial function and cell viability are interconnected with calcium signaling. Shown in Figure 6A and B, expression of Bnip3-FL reduced steady-state ER calcium levels. Surprisingly, Bnip3ΔExon3 also reduced ER calcium content and no cumulative effect on calcium release was observed with co-expression of both splice variants. Next, we examined if mitochondria were the destination of the released calcium using the mitochondrial-targeted GECO (Figure 6C, D). We observed that Bnip3-FL increased mitochondrial calcium levels. Whereas Bnip3ΔExon3 had no effect when acting alone, but when together with Bnip3-FL, Bnip3ΔExon3 attenuated mitochondrial calcium accumulation. A similar phenomenon was observed with BNIP3ΔExon2 (Figure 6E). Likewise, misoprostol had a similar effect on mitochondrial calcium accumulation when co-treating with CoCl_2_, to induce Bnip3-FL expression (Figure 6F). Moreover, knock-down of Bnip3-FL with an siRNA prevented CoCl_2_-induced mitochondrial calcium accumulation (Figure 6G). In concert with our earlier observations, knockdown of Bnip3ΔExon3 reduced the effectiveness of misoprostol in moderating the mitochondrial calcium load (Figure 6H). These observations led us to hypothesize that Bnip3ΔExon3 attenuates mitochondrial permeability transition by blocking mitochondrial calcium uptake, but not the release of calcium from the ER. To test this hypothesis, we expressed Bnip3-FL, with and without the inositol triphosphate receptor (IP3R) inhibitor 2-APB, and evaluated mitochondrial calcium levels. Shown in Figure 6I, Bnip3-FL-induced mitochondrial calcium accumulation was attenuated by 2-APB, suggesting that the ER calcium release is buffered by the mitochondria. Next, we evaluated whether pharmacological inhibition of the mitochondrial calcium uniporter (MCU) with Ru360, or inhibition of the mitochondrial voltage-dependent anion channel (VDAC) with DIDS, could affect Bnip3-FL-induced mitochondrial calcium accumulation. We observed that both Ru360 and DIDS prevented mitochondrial calcium accumulation caused by Bnip3-FL (Figure 6J). Interestingly, a combination of both inhibitors did not produce any additional effect, and that levels of mitochondrial calcium when treated with the inhibitors was similar in magnitude to those observed with co-expression of Bnip3ΔExon3. Finally, we used the SERCA inhibitor thapsigargin (Thaps) to circumvent a regulated ER calcium release. Shown in Figure 6K, thapsigargin treatment led to mitochondrial calcium accumulation that was attenuated by expression of Bnip3ΔExon3, indicating that Bnip3ΔExon3 affects entry of calcium into the mitochondria. Collectively, these observations support our hypothesis that Bnip3-FL promotes ER calcium release that accumulates in the mitochondrial matrix, while Bnip3ΔExon3 and BNIP3ΔExon2 serve to block mitochondrial calcium accumulation.

**Figure 6.**
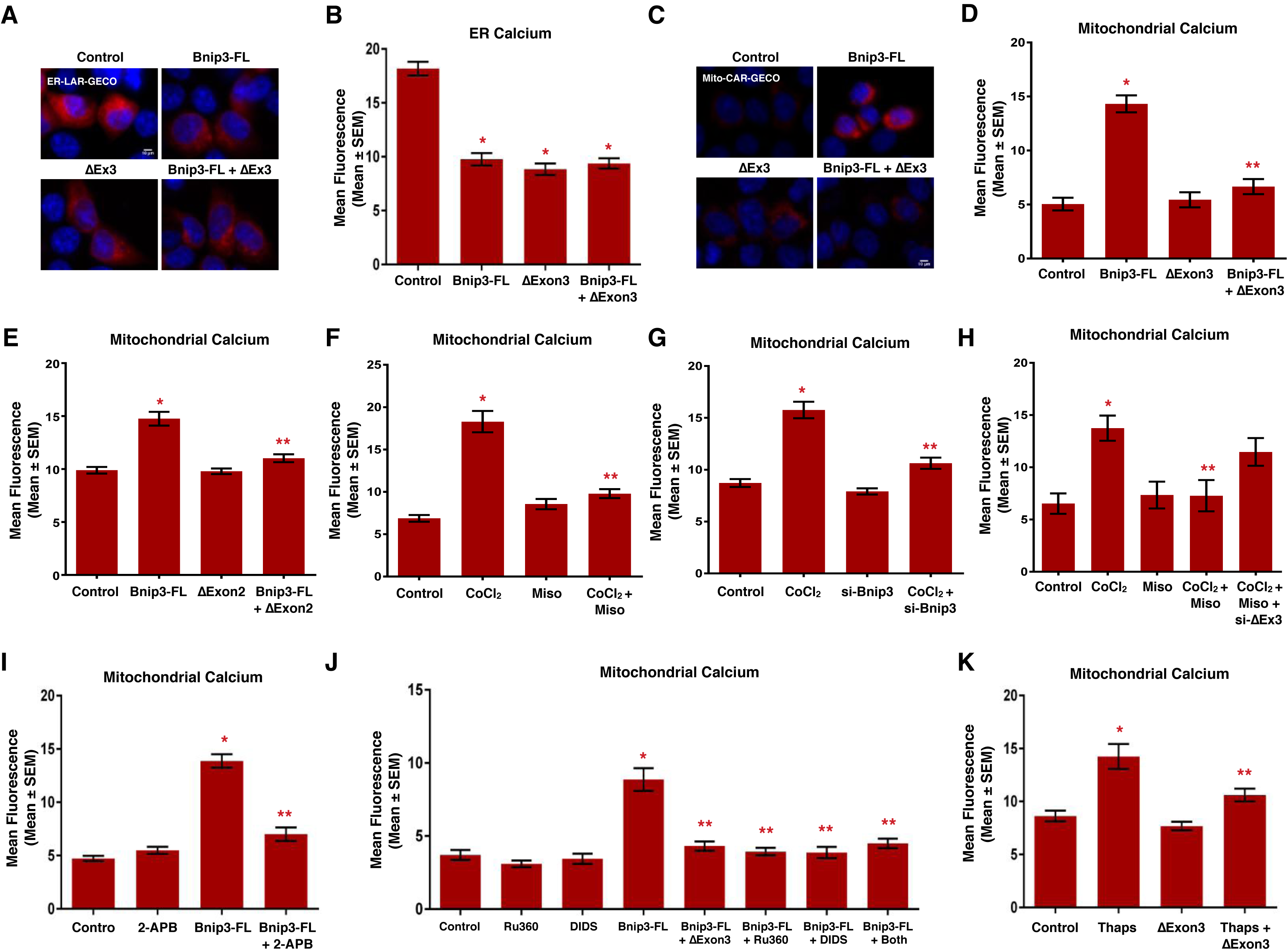
Bnip3 splice variants differentially regulate subcellular calcium micro-domains. (**A**) HCT-116 cells were transfected with Bnip3-FL, Bnip3ΔExon3 or an empty vector control. ER-LAR-GECO (red) was used to indicate endoplasmic reticulum calcium content in all conditions. Cells were stained with Hoechst (blue) and imaged by standard fluorescence microscopy. (**B**) Quantification of (A), where red fluorescent signal was normalized to cell area and quantified in 10 random fields. (**C**) HCT-116 cells were transfected with Bnip3-FL, Bnip3ΔExon3 or an empty vector control. Mito-CAR-GECO (red) was used to indicate mitochondrial calcium content in all conditions. Cells were stained and imaged as in (A). (**D**) Quantification of (C). (**E**) Quantification of HCT-116 cells transfected with Mito-CAR-GECO, Bnip3-FL, BNIP3ΔExon2 or an empty vector control. (**F**) Quantification of HCT-116 cells transfected with Mito-CAR-GECO and treated with 200 μM cobalt chloride ± 10 μM misoprostol or vehicle control for 20hrs. (**G**) Quantification of HCT-116 cells transfected with Mito-CAR-GECO, si-BNIP3-FL or scramble control. Cells were treated with 200 μM cobalt chloride or vehicle control for 16hrs. (**H**) Quantification of H9c2 cells treated with 200 μM cobalt chloride ± 10 μM misoprostol or vehicle control for 20hrs, and transfected with Mito-CAR-GECO and si-Bnip3ΔExon3 or a scrambled control. (**I**) Quantification of HCT-116 cells transfected with Mito-CAR-GECO, Bnip3-FL ± 2 μM 2-APB for 16hrs. (**J**) Quantification of HCT-116 cells treated with 10 μM DIDs and/or 10 μM Ru360 for 16hrs (where *both* indicates DIDs + Ru360) and transfected Mito-CAR-GECO, Bnip3-FL, Bnip3ΔExon3 and/or empty vector control. (**K**) Quantification of HCT-116 cells treated with 1 μM Thapsigargin (Thaps) for 4hrs and transfected with Mito-CAR-GECO, Bnip3ΔExon3 or empty vector control. Data are represented as mean ± S.E.M. *^*^P*<0.05 compared with control, while *^**^P*<0.05 compared with treatment, determined by 1-way ANOVA.

Bnip3 was originally discovered as an interacting partner of the pro-survival protein BCL-2^37^. Interestingly, BCL-2 has been implicated as an important regulator of cellular calcium signaling through multiple mechanisms, including blocking mitochondrial calcium uptake through inhibition of VDAC channels^34,38^. Thus, we investigated the possibility of BCL-2 involvement in Bnip3ΔExon3-regulated calcium signaling. Specifically, we tested if Bnip3ΔExon3 could alter the cellular location of BCL-2. We expressed Flag-tagged BCL-2, with and without HA-tagged Bnip3ΔExon3, in HCT-116 cells and performed mitochondrial and cytosolic fractionation. Western blot analysis of these fractions revealed that Bnip3ΔExon3 reduced the cytosolic levels of BCL-2 and increased the mitochondrial BCL-2 levels, without impacting the whole-cell expression of BCL-2 (Figure 7A). These findings suggest that Bnip3ΔExon3 promotes mitochondrial translocation of BCL-2. Additionally, we observed that both BCL-2 and the structurally related protein MCL-1, were capable of blocking Bnip3-FL-induced mitochondrial calcium accumulation (Figure 7B).

**Figure 7.**
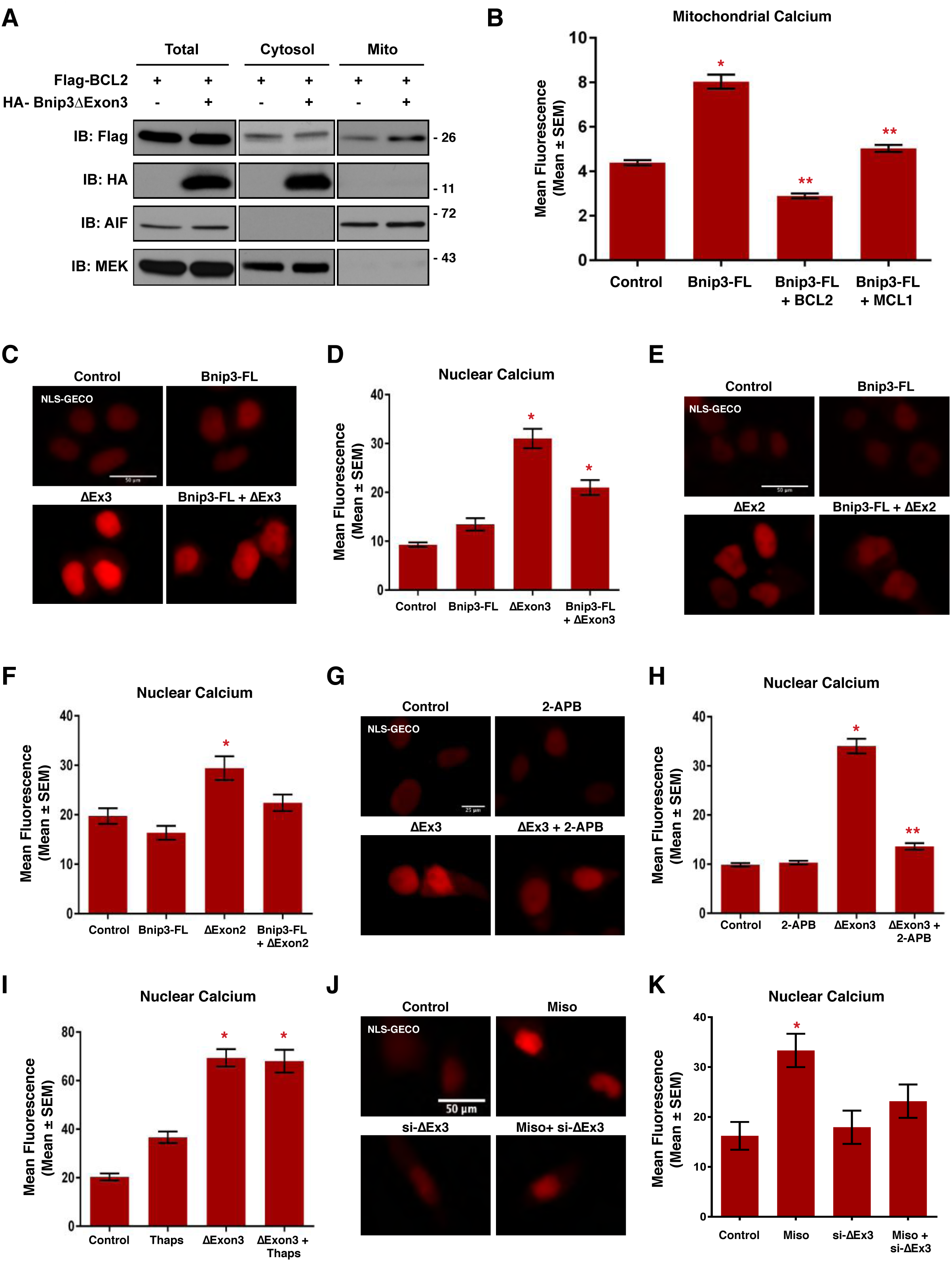
Bnip3 splice variants regulate nuclear calcium content. (**A**) HCT-116 cells were transfected with Flag-BCL2 and HA-Bnip3ΔExon3, as indicated. Protein extracts were subjected to fractionation and were immunoblotted, as indicated. (**B**) HCT-116 cells were transfected with Mito-CAR-GECO, Bnip3-FL, Flag-BCL2, or MCL-1, as indicated. Fluorescence was normalized to cell area and quantified in 10 random fields. (**C**) HCT-116 cells were transfected with Bnip3-FL, Bnip3ΔExon3 or an empty vector control. NLS-R-GECO (red) was used to indicate nuclear calcium content. Cells were stained with Hoechst (blue) and imaged by standard fluorescence microscopy. (**D**) Quantification of (C), where red fluorescent signal was normalized to nuclear area and quantified in 10 random fields. (**E**) HCT-116 cells were transfected with NLS-R-GECO, Bnip3-FL, BNIP3ΔExon2 or an empty vector control. (**F**) Quantification of (E). (**G**) HCT-116 cells were transfected with NLS-R-GECO, Bnip3ΔExon3 ± 2 μM 2-APB for 16hrs. (**H**) Quantification of (G). (**I**) HCT-116 cells transfected with NLS-R-GECO, Bnip3ΔExon3 ± 1 μM Thapsigargin (Thaps) for 4hrs. (**J**) H9c2 cells transfected NLS-R-GECO, si-Bnip3ΔExon3 (indicated as si-ΔEx3) or scrambled control, and treated with 10 μM misoprostol or vehicle control for 20hrs. Cells were stained with Hoechst (blue) and imaged by standard fluorescence microscopy. (**K**) Quantification of (J). Data are represented as mean ± S.E.M. *^*^P*<0.05 compared with control, while *^**^P*<0.05 compared with treatment, determined by 1-way ANOVA or unpaired t-test.

Our observation that Bnip3ΔExon3 did not block ER calcium release, but did prevent mitochondrial calcium accumulation, led us to investigate the nucleus as a possible subcellular destination for this calcium. Using a GECO-based calcium biosensor fused to a nuclear localization signal (NLS-R-GECO)^35^, we evaluated the effect of Bnip3ΔExon3 on nuclear calcium content. Shown in Figure 7C and D, Bnip3ΔExon3 increased nuclear calcium levels, whereas Bnip3-FL had no effect. Although not as marked of an elevation as observed with Bnip3ΔExon3 alone, nuclear calcium remained elevated when Bnip3-FL was co-expressed with Bnip3ΔExon3. Similar effects on nuclear calcium were observed with BNIP3ΔExon2 (Figure 7E, F). In addition, we observed that Bnip3ΔExon3-induced calcium accumulation was prevented by the IP3R inhibitor 2-APB (Figure 7G, H), providing evidence for a dependence on ER-calcium as the source of nuclear calcium. Once again, we used thapsigargin to circumvent a regulated ER calcium release and observed no additional effect on Bnip3ΔExon3-induced nuclear calcium accumulation, indicating that this is an active process mediated by Bnip3ΔExon3 (Figure 7I). Finally, we examined the interconnectivity of misoprostol and Bnip3ΔExon3 in this signaling pathway. We found that treatment with misoprostol also increased nuclear calcium levels. Importantly, this effect was diminished by siRNA knockdown of Bnip3ΔExon3 (Figure 7J, K). Collectively, these observations provide evidence that Bnip3ΔExon3 promotes ER-to-nuclear calcium transfer, while Bnip3-FL promotes ER-to-mitochondrial calcium transfer. Furthermore, mitochondrial translocation of BCL-2 might be an important intermediate step in the distinct regulation of calcium signals initiated by Bnip3ΔExon3.

### Bnip3ΔExon3 promotes cardiomyocyte hypertrophy

In order to understand the cellular consequences of Bnip3ΔExon3-induced nuclear calcium accumulation, we evaluated two transcriptional regulators known to be regulated in a calcium-dependent manner. Theoretically, these regulators (NFATc3 and HDAC5) offer a tool to test the potential of Bnip3ΔExon3-regulated calcium signalling as a modulator of gene expression; biologically, these tools serve as indications of the cellular significance of this regulatory pathway. First, the transcription factor NFATc3 is activated by the calcium-calmodulin activated phosphatase calcineurin, promoting nuclear translocation. Second, the class II histone deacetylase, HDAC5, is exported from the nucleus upon phosphorylation by calcium-calmodulin activated kinases (CaMKs). Thus, we evaluated the effect of Bnip3ΔExon3 on the subcellular localization of these proteins as an indication of their activity by way of fluorescent fusion proteins of NFATc3 (NFAT-YFP) and HDAC5 (HDAC5-GFP). We observed that expression of Bnip3ΔExon3 increased the nuclear localization of NFAT-YFP (Figure 8A, B), and increased the cytosolic localization of HDAC5-GFP (Figure 8C, D). These observations suggest that Bnip3ΔExon3-induced nuclear calcium has the potential to alter gene expression. Interestingly, both NFATs and class II HDACs have been implicated as regulators of cardiomyocyte growth during development and cardiac disease. Thus, as a test of the cellular significance of this regulatory axis we transduced primary neonatal cardiomyocytes with a lentivirus delivering Bnip3ΔExon3 and performed immunofluorescence with the sarcomeric myosin antibody MF-20 to measure cell area. Shown in Figure 8E and F, we observed a statistically significant increase in cardiomyocyte cell area with Bnip3ΔExon3 expression. Furthermore, we evaluated the expression of two classical marker genes of cardiac hypertrophy and reactivation of the ‘fetal gene program’, α- and β-myosin heavy chain (MHC), by real-time PCR. In this regard, a shift from α- MHC (*Myh6*) to β-MHC (*Myh7*) serves as an indication of the reversion from ‘adult’ to ‘fetal’ gene expression programs^39,40^. We observed that Bnip3ΔExon3 increased *Myh7* expression and reduced *Myh6* expression (Figure 8G). This resulted in an increase in the MHC ratio (*Myh7*/*Myh6*; 1.66) with Bnip3ΔExon3 transduction. Similarly, we observed that treatment of primary cardiomyocytes with misoprostol also increased cell area, and the magnitude of this effect was matched in cells treated with the well-established hypertrophy-inducer phenylephrine (PE) (Figure 8H). Collectively, these findings suggest that nuclear accumulation of calcium elicited by the small Bnip3 splice variants operates to regulate the activity of transcription regulators and has the potential to control cardiomyocyte growth, in addition to their roles in cell survival.

**Figure 8.**
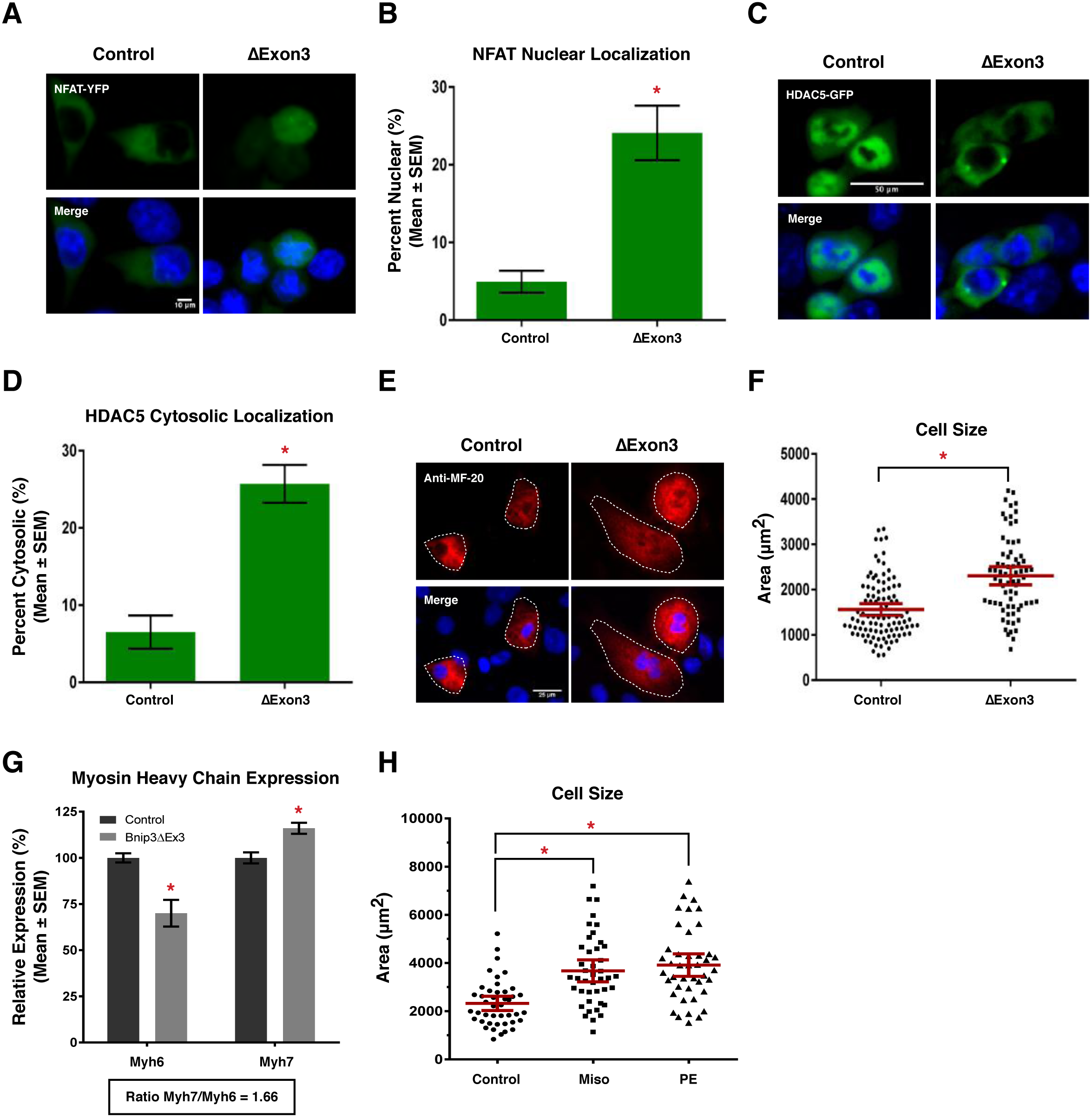
Bnip3ΔExon3 regulates cardiomyocyte hypertrophy. (**A**) HCT-116 cells were transfected with Bnip3ΔExon3 or an empty vector control. NFAT-YFP (green) was used to indicate subcellular localization of NFAT. Cells were stained with Hoechst (blue) and imaged by standard fluorescence microscopy. (**B**) Quantification of fluorescent images in (A), by calculating the percentage of cells with nuclear NFAT signal over 10 random fields. (**C**) HCT-116 cells were transfected with Bnip3ΔExon3 or an empty vector control. HDAC5-GFP (green) was used to indicate subcellular localization of HDAC5. Cells were stained and imaged as in (A). (**D**) Quantification of fluorescent images in (C), by calculating the percentage of cells with cytosolic HDAC5 signal over 10 random fields. (**E**) Primary ventricular neonatal cardiomyocytes (PVNM) cells were transduced with Bnip3ΔExon3 or control virus. Cells were fixed, stained with Hoechst (blue), and probed for myosin heavy chain (Anti-MF-20, red) expression. (**F**) Quantification of (E), where cell size (μm^2^) was calculated based on the area of the red fluorescent signal and quantified in 10 random fields. (**G**) PVNM cells treated as in (E). RNA was isolated and qRT-PCR was performed for myosin heavy chain isoform expression. (**H**) Quantification of PVNM cells treated with 10 μM misoprostol, 10 μM phenylephrine (PE) or vehicle control for 20hrs. Cells were stained with calcein-AM (green) and assessed for cell size. Cell size (μm^2^) was calculated based on the area of the green fluorescent signal and quantified in 10 random fields. Data are represented as mean ± S.E.M. *^*^P*<0.05 compared with control, determined by unpaired t-test.

## Discussion

The cellular response to hypoxia involves both adaptive cell survival phenomenon and regulated cell death; however, the mechanisms dictating these opposing cell phenotypes during development and disease remain poorly defined. Here we provide evidence that activation of prostaglandin signaling through misoprostol treatment induces pro-survival NF-κB gene expression during hypoxia, resulting in reduced expression of the pro-death Bnip3-FL splice variant and increased expression of pro-survival small Bnip3 variants (Figure 9A). Our data also implicates the combined action of HIF1α and NF-κB P65 to promote cell survival, whereas activation of HIF1α alone induces Bnip3-FL expression and cell death. We also demonstrate that the *BNIP3* gene can be differentially spliced in human versus rodent cells.

**Figure 9.**
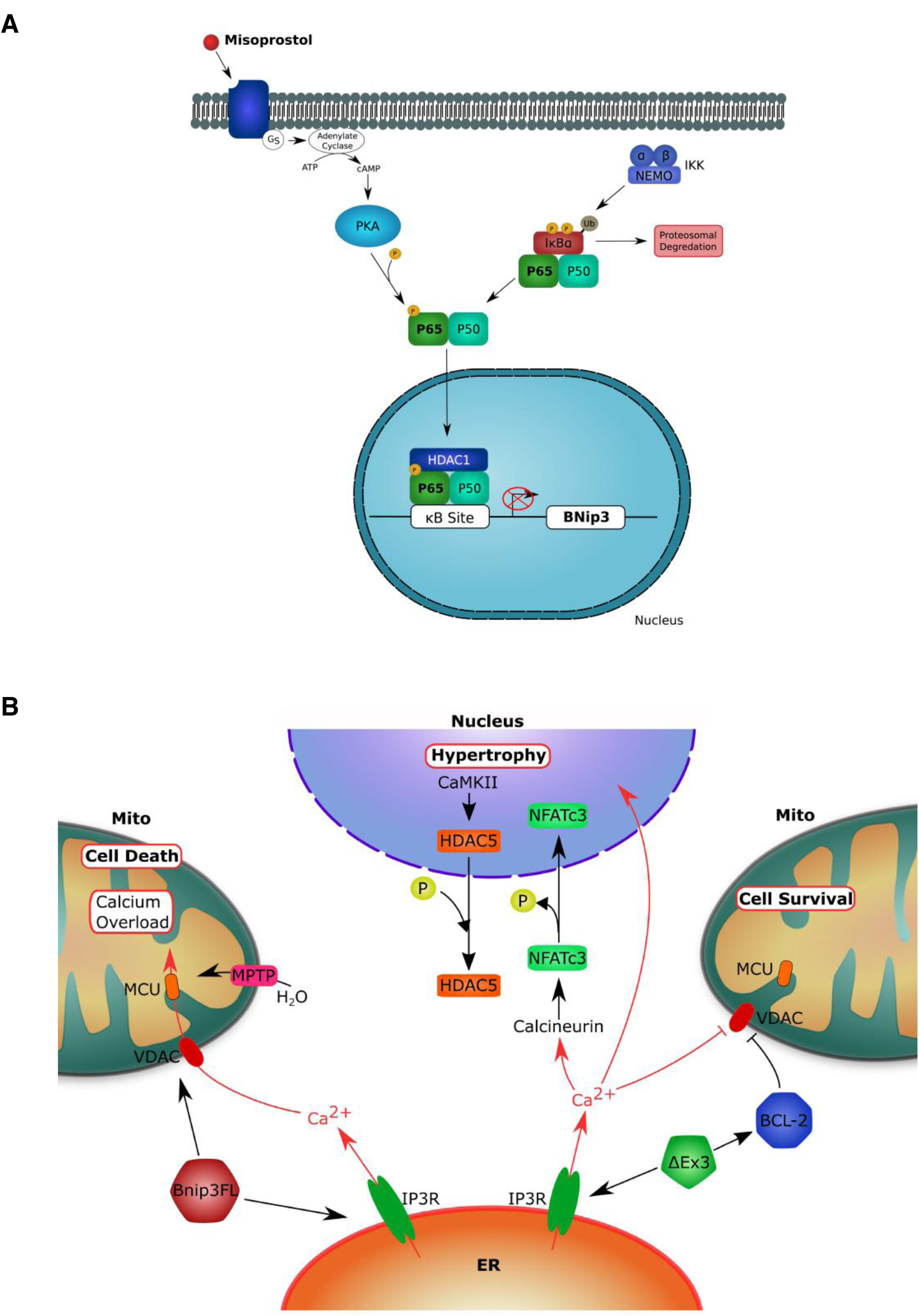
Proposed mechanisms of misoprostol-induced cellular protection. (**A**) Proposed mechanism by which misoprostol opposes Bnip3-FL expression. (**B**) Proposed mechanism by which Bnip3ΔExon3 regulates calcium localization, cell survival and hypertrophy.

Since its discovery, it has been noted that Bnip3 could induce context-dependent apoptotic or necrotic cell death^14,37^. Early studies evaluating Bnip3 determined its subcellular location is predominantly mitochondrial, and cells expressing Bnip3 displayed features consistent with apoptosis^37^. However, more detailed analysis of Bnip3 revealed that it could activate cell death from extra-mitochondrial sites and cells expressing Bnip3 display a necrotic cell phenotype, particularly when Bnip3 is targeted to the ER with the cytochrome-b5 transmembrane domain^12,14^. Importantly, cell death elicited by Bnip3 is attenuated with the mitochondrial permeability transition pore inhibitor cyclosporine-A and the mitochondrial calcium uniporter blocker Ru360^12,14^; whereas, cells expressing BNIP3 are resistant to caspases inhibitors, such as zVAD-fmk^14^. These findings are consistent with the notion that Bnip3 is an important regulator of mitochondrial permeability transition-regulated necrosis, through a calcium-dependent mechanism^28,29^. Current models of Bnip3 function suggest that Bnip3 regulates apoptosis and mitophagy when localized at the mitochondria, and macro-autophagy and necrosis when localized at the ER membranes^13^. However, a definitive mechanism describing how Bnip3 is translocated to either the mitochondria or ER is lacking, and many questions remain regarding cell-type specific differences in Bnip3 function.

The results of the present study expand these previous observations and suggest that alternative splicing of Bnip3 is an important mechanism to control cellular calcium homeostasis, cell death, and cell growth. We observed that both Bnip3-FL and Bnip3ΔExon3 deplete ER calcium content. However, Bnip3-FL preferentially transfers calcium to the mitochondrial matrix and triggers mitochondrial permeability transition and necrosis; whereas, Bnip3ΔExon3 serves to block mitochondrial accumulation and promote nuclear calcium transfer leading myocyte cell growth (Figure 9B). However, the role of the endogenous Bnip3 splice variants has not been fully validated *in vivo*, and the function of specific splice variants has yet to be characterized by gene-targeting studies. Nonetheless, these findings serve to unify the seemingly disparate observations of others who have observed that IP3R-dependent calcium release can induce cell death^33,34^, and cardiac hypertrophy^41^. Our findings suggest that the splicing of Bnip3, and the cellular location of BCL-2, can influence cell fate by dictating where the IP3R-dependent calcium release accumulates, either in the mitochondria or the nucleus. However, additional *in vivo* experimentation is required to evaluate the role of Bnip3 splicing in hypertrophic regulation and determine under which biological contexts this Bnip3-dependent mechanism operates, as other BCL-2 family members, such as Bnip3L/Nix, likely contribute to cardiac calcium homeostasis^36^.

Another important observation in the present study is the identification of the human BNIP3ΔExon2 splice variant. Although we identified several similarities between BNIP3ΔExon2 and Bnip3ΔExon3, such as inhibition of cell death, prevention of ER-mitochondrial calcium transfer, and nuclear calcium accumulation, potential differences likely exist. *In silico* domain mapping revealed conserved N-terminal and C-terminal sequences; however, significant sequence differences exist, including a predicted nuclear localization sequence present in BNIP3ΔExon2 that is not present in Bnip3ΔExon3, and the LIR domain which is present in Bnip3ΔExon3 but not BNIP3ΔExon2. Further experimental work is required to fully define how BNIP3ΔExon2 and Bnip3ΔExon3 functions are distinct. Interestingly, recent work regarding the Bnip3 homologue, Bnip3L/Nix, has demonstrated that its small splice variant, named sNix, alters gene expression to promote cell survival in an NF-κB-dependent manner^42^. Collectively, this suggests an interplay between Bnip3 and Bnip3L/Nix splicing and NF-κB-dependent survival gene expression. Moreover, the expression ratio of Nix to sNix was determined to be approximately 10:1, which is consistent with our findings regarding ratio of Bnip3 to its small survival splice variants, and suggests that these small splice variants exert their physiological effects through gene expression amplification rather than direct inhibition of the full-length isoforms^42^.

Recent progress has been made regarding the mitochondrial permeability transition pore structure and function, and several studies have defined the importance of permeability transition-dependent necrosis in the pathophysiology of ischemic cardiac and cerebrovascular diseases^29^. Our data suggests that Bnip3 splicing plays an important role in the complications arising from neonatal hypoxia. However, additional studies are needed to determine if this mechanism operates to control permeability transition, necrosis, and/or cell growth in other pathologies. Given the diverse roles of Bnip3 splice variants regulating mitochondrial function, autophagy, and cell growth, dysregulation of this genetic pathway may have broad implications involving ischemic cardiovascular and cerebrovascular diseases, as well as cancer biology.

## Materials and Methods

### In vivo neonatal hypoxia model

All procedures in this study were approved by the Animal Care Committee of the University of Manitoba, which adheres to the principles for biomedical research involving animals developed by the Canadian Council on Animal Care. Litters of Long-Evans rat pups (N=5) and their dam were placed into a hypoxia chamber with 10% oxygen delivery, at post-natal day-3 (PND3) for 7 days. Control pups (N=5) were left in normoxic conditions at 21% oxygen^23,24^. Animals received subcutaneous injections of 10 μg/kg misoprostol, or saline control, for 7 days. At PND10 animals were euthanized and perfused with saline for tissue collection.

### Plasmids and virus production

The PKA biosensor (pPHT-PKA) was a gift from Anne Marie Quinn (Addgene #60936)^25^. The plasmids for human P65 expression including: GFP-RelA, T7-RelA, T7-RelA(S276A), and PCMV4-3 HA/IkB-alpha(SS32,36AA) were gifts from Warner Greene (Addgene #23255, 21984, 24153, 24143)^43,44^. The plasmid for HIF1α (HA-HIF1α-pcDNA3) was a gift from William Kaelin (Addgene #18949)^45^. The mitochondrial (CMV-mito-CAR-GECO1), endoplasmic reticulum (CMV-ER-LAR-GECO1), and nuclear (CMV-NLS-R-GECO) targeted calcium biosensors were gifts from Robert Campbell (Addgene #46022, 61244, and 32462)^35,46,47^. Flag-BCL-2 was gift from Clark Distelhorst (Addgene #18003)^48^. pMSCV-puro-mMCL1-WT was a gift from Joseph Opferman (Addgene #45817)^49^. NFATc3 fused to YFP (NFAT-YFP) was a gift from Tetsuaki Miyake, and HDAC5-GFP was provided by E. Olson. The mouse myc-Bnip3-FL and HA-Bnip3ΔExon3 (Accession #MF156210) were described previously (Addgene #100796, #100793)^22^. The pLenti-Bnip3ΔExon3 virus was generated by ligating a PCR amplicon (HA-Bnip3ΔExon3) into the pLenti-puro back bone, which was a gift from Ie-Ming Shih (Addgene #39481)^50^. Virus was packaged and purified in the University of Manitoba Lentiviral Vector Viral Particles & Production Core. Human HA-BNIP3-FL and HA-BNIP3ΔExon2 (Accession #MF593120) (Addgene #100781, #100782) were generated by PCR using cDNA from HUECs and primers containing EcoRI and XhoI restriction enzymes sites in the forward and reverse primers, respectively. Primers include BNIP3 Forward: 5’-TATGCG<u>GAATTC</u>ATGTCGCAGAACGGAGCGCCCGGGAT-3’, BNIP3 Reverse: 5’-AATCCG<u>CTCGAG</u>TCAAAAGGTGCTGGTGGAGGTTGTC-3’; BNIP3ΔExon2 Reverse: 5’-AATCCG<u>CTCGAG</u>TCAAGATGCTTTCAACTTCTTTCC-3’. PCR products were cut and ligated into pcDNA3, in frame with an N-terminal HA tag, described previously^51,52^.

### Cell culture, transductions and transfections

HCT-116 cells were maintained in McCoy’s 5A medium (Hyclone), containing penicillin, streptomycin, and 10% fetal bovine serum (Hyclone), at 37°C and 5% CO_2_. H9c2 and 3T3 cells were maintained in Dulbecco’s modified Eagle’s medium (DMEM; Hyclone), containing penicillin, streptomycin, and 10% fetal bovine serum (Hyclone), at 37°C and 5% CO_2_. Rat primary ventricular neonatal cardiomyocytes (PVNC) were isolated from 1-2 day old pups using the Pierce Primary Cardiomyocyte Isolation Kit (#88281). All cells were transfected using JetPrime Polyplus reagent, as per the manufacturer’s protocol^30^. Overexpression of Bnip3ΔExon3 in PVNC cells was carried out using pLenti-puro-Bnip3ΔExon3 lentiviral particles, alongside control lentiviral particles (Santa Cruz, sc-108080) to control for the effects of transduction alone. RNAi experiments targeting Bnip3-FL and Bnip3ΔExon3 were performed using siRNA targeting the 3^rd^ exon (sense sequence: UCGCAGACACCACAAGAUA) and the exon 2-4 junction (5’-CACUGUGACAGUCUGAGGA) respectively (Dharmacon). A scrambled siRNA was used as control (Santa Cruz sc-37007). siRNA’s were transfected into cells using JetPrime Polyplus reagent, as per the manufacturer’s protocol^30^. For misoprostol treatments, 10mM misoprostol (Sigma) in phosphate buffered saline (PBS; Hyclone) was diluted to 10μM directly in media and applied to cells for 20 hours^22^. For H89 treatment, a 10mM stock of H89-dyhydrochloride hydrate (Sigma) in PBS, was diluted to 10μM directly in media. For cobalt chloride (CoCl_2_) treatment, a 200mM stock solution of cobalt(II) chloride hexahydrate (Sigma) in PBS, was diluted to 200μM directly in media and applied to cells. CoCl_2_ concentration was selected based on previously published data looking at HIF1α induction^53^. For calcium imaging experiments, 1μM Thapsigargin (Sigma) was applied directly to cells for 4 hours, while 2 μM 2-APB (Sigma), 10μM Ru360 (Sigma), and 10μM disodium 4,4′-diisothiocyanatostilbene-2,2′-disulfonate (DIDS, Sigma) were applied directly to cells for 16 hours. For cardiac hypertrophy experiments, a 10mM stock of phenylephrine (Sigma) in PBS, was diluted to 10μM directly in media.

### Fluorescent staining and live cell imaging

Hoechst 33342, TMRM, calcein-AM and Ethidium Homodimer-1 were all purchased from Biotium and applied using manufacturer’s protocol^30,54^. MitoSOX was purchased from Life Technologies and was applied to cells for a 10-minute incubation to assess mitochondrial superoxide^30^. Calcein-Cobalt Chloride (CoCl_2_) staining was used to assess mitochondrial permeability transition, which is achieved through quenching cytosolic calcien-AM signal with 5μM CoCl_2_ described previously^30^. Immunofluorescence with Anti-MF-20 (DSHB # AB_2147781) was used with fluorescent secondary antibody conjugated to Alexa Fluor 555 (Themo # A-31570) to assess cardiac hypertrophy in fixed and permeabilized PVNM cells. All imaging experiments were done on an Olympus IX70 inverted microscope with QImaging Retiga SRV Fast 1394 camera using NIS Elements AR 3.0 software. Quantification, scale bars, and processing including background subtraction, was performed on Fiji (ImageJ) software.

### Immunoblotting

Protein extractions were achieved using a RIPA lysis buffer combined with Phenylmethanesulfonyl fluoride (PMSF, Sigma), Sodium orthovanadate (NaVO, Sigma) and phosphatase inhibitor cocktail (PIC, Santa Cruz). Homogenization was required in the case of tissue protein extraction. Mitochondrial/cytosolic fractionation was done using a Mitochondrial Isolation Kit (Qiagen Qproteome # 37612), while nuclear/cytosolic fractionation was done using a NE-PER Nuclear and Cytosolic Extraction Kit (Pierce # 78833), as described previously^21,55,56^. Protein content was determined using a Bio-Rad Protein Assay Kit. Extracts were resolved via SDS-PAGE and later transferred to a PVDF membrane using an overnight transfer system. Immunoblotting was carried out using primary antibodies in 5% powdered milk or BSA (as per manufacturer’s instructions) dissolved in TBST. Horseradish peroxidase-conjugated secondary antibodies (Jackson ImmunoResearch Laboratories; 1:5000) were used in combination with enhanced chemiluminescence (ECL) to visualize bands^55,56^. The following antibodies were used: Bnip3-FL and BNIP3ΔExon2 (CST # 44060), P65 (CST # 8242), P65 Phospho-Ser276 (Assay BioTech # A7169), HIF1α (CST # 14179), Flag (CST # 2368), HA (CST # 3724), AIF (CST # 5318), MEK (CST # 8727), Histone-H3 (CST # 4499), Tubulin (CST # 86298), and Actin (Santa Cruz sc-1616). For the detection of rodent Bnip3 splice variants a custom rabbit polyclonal antibody was generated by Abgent using the following peptide sequence CSQSGEENLQGSWVE.

### Mitochondrial respiration

Mitochondrial respiration in PVNM cells was assessed using the Seahorse XF-24 Extracellular Flux Analyzer (Seahorse Bioscience, North Billerica, MA, USA), as described previously^30^. Calculated respiration rates were determined as per manufacturer’s instructions (Mito Stress Kit; Seahorse Bioscience).

### Reverse Transcription PCR

Total RNA was extracted from cells and pulverized tissues by the TRIzol extraction method and genomic DNA was removed via the RNeasy Mini Kit (Qiagen), including an On-Colum DNase I Digestion (Qiagen). For reverse transcription PCR (RT-PCR), purified mRNA was reverse transcribed using Maxima Enzyme (Thermo), followed by PCR using Taq polymerase (New England Biolabs). Amplified RNA was then run on a 2% agarose gel with GelRed nucleic acid gel stain (Biotium). The cloning primers, described above, were used to detect human BNIP3 splice variants. Mouse Bnip3 primers were: Forward: 5’- GCCG<u>GAATTC</u>ATGTCGCAGAGCGGGGAG-3’, Bnip3-FL Reverse: 5’- CGGCG<u>CTCGAG</u>TCAGAAGGTGCTAGTGGAAGTTGTC-3’, and Bnip3ΔExon3 Reverse: 5’- CGGCG<u>CTCGAG</u>TCAGGATACTTTCAACTTCTCTTCTTCTCTC-3’ Primers used for rat heart tissue were: 5’ TTCCAGCTTCCGTCTCTATTT-3’ and 5’- TCAGGATACTTTCAACTTCTCTTCT-3’. For quantitative real-time PCR (qPCR) targeting myosin heavy chain expression, mRNA was extracted from cells using TRIzol then reverse transcribed into cDNA. Following DNase treatment, cDNA was combined with SYBR Green Supermix (Thermo) and mRNA was amplified using the ABI 7500 Real-Time PCR system (Applied Biosystems). Primers were: Rat myosin heavy chain 6 forward: 5’- GAGGAATAACCTGTCCAGCAG -3’, and reverse: 5’-TACAGGCAAAGTCAAGCATTC -3’. Rat myosin heavy chain 7 forward: 5’- CCAACACCAACCTGTCCAAG-3’, and reverse: 5’- CAAAGGCTCCAGGTCTCAGG -3’. Rat B-Actin forward: 5’- CTGTGTGGATTGGTGGCTCTA -3’, and reverse 5’- AAAACGCAGCTCAGTAACAGTCC -3’.

### Statistics

Data are presented as mean ± standard error of the mean (S.E.M.). Differences between groups in imaging experiments with only 2 conditions were analyzed using an unpaired t-tests, where (*) indicates *P*<0.05 compared with control. In the case of the cell size experiment, a 2-tailed t-test with Welches correction was used to compare the differences between conditions. Experiments with 4 or more conditions were analyzed using a 1-way ANOVA, with Tukey’s test for multiple comparison, where (*) indicates P<0.05 compared with control, and (**) indicates P<0.05 compared with treatment. All statistical analysis was done using GraphPad Prism 6 software.

## Acknowledgements

This work was support by the Natural Science and Engineering Research Council (NSERC) Canada, through a Discovery Grant to JWG and a studentship to JF. Seed funding was provided by the Children’s Hospital Foundation of Manitoba. WDJ is funded by a RIG grant from Athabasca University and TI is funded through the SFRG program from the University of Manitoba. GMH and JWG are supported by the Heart and Stroke Foundation of Canada and are members of the DEVOTION Research Cluster. GMH is a Canada Research Chair in Molecular Cardiolipin Metabolism. WM is supported by a scholarship from the Children’s Hospital Foundation of Manitoba and Research Manitoba, MM received support from the DEVOTION research cluster. We wish to thank Dr. Tetsuaki Miyake for the NFAT-YFP plasmid and Caitlin Blaney for assistance the with neonatal hypoxia model and misoprostol injections.

## Conflicts

None.

## Author contributions

JWG, WDJ, and TLI conceived and coordinated the study. JWG, WDJ, TLI, MDM, and JTF wrote the paper. JTF and MDM designed and conducted most of the experiments. JTF and MDM were also responsible for data analysis and presentation. WM designed and conducted mitochondrial respiration experiments. YH designed and conducted the RT-PCR experiments, and designed and constructed all novel plasmids. JWG performed *in silico* work, including the 3D modeling and alignments. TLI and MDM designed and conducted the in vivo hypoxia experiments. JWG, WDJ, TLI and GMH were responsible for funding acquisition. All authors reviewed the results, edited, and approved the final version of the manuscript.

## References

1 Douglas-Escobar M, Weiss MD. Hypoxic-ischemic encephalopathy: a review for the clinician. JAMA Pediatr 2015; 169: 397–403.

2 Neu J, Walker WA. Necrotizing enterocolitis. N Engl J Med 2011; 364: 255–264.

3 Wang H, Zhang SX, Hartnett ME. Signaling pathways triggered by oxidative stress that mediate features of severe retinopathy of prematurity. JAMA Ophthalmol 2013; 131: 80–85.

4 Dakshinamurti S. Pathophysiologic mechanisms of persistent pulmonary hypertension of the newborn. Pediatr Pulmonol 2005; 39: 492–503.

5 Armstrong K, Franklin O, Sweetman D, Molloy EJ. Cardiovascular dysfunction in infants with neonatal encephalopathy. Arch Dis Child 2012; 97: 372–375.

6 Shastri AT, Samarasekara S, Muniraman H, Clarke P. Cardiac troponin I concentrations in neonates with hypoxic-ischaemic encephalopathy. Acta Paediatr Oslo Nor 1992 2012; 101: 26–29.

7 Greer SN, Metcalf JL, Wang Y, Ohh M. The updated biology of hypoxia-inducible factor. EMBO J 2012; 31: 2448–2460.

8 Carmeliet P, Dor Y, Herbert JM, Fukumura D, Brusselmans K, Dewerchin M et al. Role of HIF-1alpha in hypoxia-mediated apoptosis, cell proliferation and tumour angiogenesis. Nature 1998; 394: 485–490.

9 Kothari S, Cizeau J, McMillan-Ward E, Israels SJ, Bailes M, Ens K et al. BNIP3 plays a role in hypoxic cell death in human epithelial cells that is inhibited by growth factors EGF and IGF. Oncogene 2003; 22: 4734–4744.

10 Azad MB, Chen Y, Henson ES, Cizeau J, McMillan-Ward E, Israels SJ et al. Hypoxia induces autophagic cell death in apoptosis-competent cells through a mechanism involving BNIP3. Autophagy 2008; 4: 195–204.

11 Mazure NM, Pouysségur J. Atypical BH3-domains of BNIP3 and BNIP3L lead to autophagy in hypoxia. Autophagy 2009; 5: 868–869.

12 Zhang L, Li L, Liu H, Borowitz JL, Isom GE. BNIP3 mediates cell death by different pathways following localization to endoplasmic reticulum and mitochondrion. FASEB J 2009; 23: 3405–3414.

13 Zhang J, Ney PA. Role of BNIP3 and NIX in cell death, autophagy, and mitophagy. Cell Death Differ 2009; 16: 939–946.

14 Vande Velde C, Cizeau J, Dubik D, Alimonti J, Brown T, Israels S et al. BNIP3 and genetic control of necrosis-like cell death through the mitochondrial permeability transition pore. Mol Cell Biol 2000; 20: 5454–5468.

15 Gordon JW, Shaw JA, Kirshenbaum LA. Multiple facets of NF-κB in the heart: to be or not to NF-κB. Circ Res 2011; 108: 1122–1132.

16 Hayden MS, Ghosh S. Shared principles in NF-kappaB signaling. Cell 2008; 132: 344–362.

17 Hayden MS, Ghosh S. Signaling to NF-kappaB. Genes Dev 2004; 18: 2195–2224.

18 Zhong H, Voll RE, Ghosh S. Phosphorylation of NF-kappa B p65 by PKA stimulates transcriptional activity by promoting a novel bivalent interaction with the coactivator CBP/p300. Mol Cell 1998; 1: 661–671.

19 Zhong H, May MJ, Jimi E, Ghosh S. The phosphorylation status of nuclear NF-kappa B determines its association with CBP/p300 or HDAC-1. Mol Cell 2002; 9: 625–636.

20 Zhong H, SuYang H, Erdjument-Bromage H, Tempst P, Ghosh S. The transcriptional activity of NF-kappaB is regulated by the IkappaB-associated PKAc subunit through a cyclic AMP-independent mechanism. Cell 1997; 89: 413–424.

21 Gang H, Dhingra R, Gordon JW, Yurkova N, Aviv Y, Aguilar F et al. A novel hypoxia-inducible spliced variant of mitochondrial death gene Bnip3 promotes survival of ventricular myocytes. Circ Res 2011; 108: 1084–1092.

22 Diehl-Jones W, Archibald A, Gordon JW, Mughal W, Hossain Z, Friel JK. Human Milk Fortification Increases Bnip3 Expression Associated With Intestinal Cell Death In Vitro. J Pediatr Gastroenterol Nutr 2015; 61: 583–590.

23 Ward NL, Moore E, Noon K, Spassil N, Keenan E, Ivanco TL et al. Cerebral angiogenic factors, angiogenesis, and physiological response to chronic hypoxia differ among four commonly used mouse strains. J Appl Physiol Bethesda Md 1985 2007; 102: 1927–1935.

24 Wallace MG, Hartle KD, Snow WM, Ward NL, Ivanco TL. Effect of hypoxia on the morphology of mouse striatal neurons. Neuroscience 2007; 147: 90–96.

25 Ding Y, Li J, Enterina JR, Shen Y, Zhang I, Tewson PH et al. Ratiometric biosensors based on dimerization-dependent fluorescent protein exchange. Nat Methods 2015; 12: 195–198.

26 Kelley LA, Mezulis S, Yates CM, Wass MN, Sternberg MJE. The Phyre2 web portal for protein modeling, prediction and analysis. Nat Protoc 2015; 10: 845–858.

27 Gordon JW. Regulation of cardiac myocyte cell death and differentiation by myocardin. Mol Cell Biochem 2017. doi:10.1007/s11010-017-3100-3.

28 Kwong JQ, Molkentin JD. Physiological and pathological roles of the mitochondrial permeability transition pore in the heart. Cell Metab 2015; 21: 206–214.

29 Izzo V, Bravo-San Pedro JM, Sica V, Kroemer G, Galluzzi L. Mitochondrial Permeability Transition: New Findings and Persisting Uncertainties. Trends Cell Biol 2016; 26: 655–667.

30 Mughal W, Nguyen L, Pustylnik S, da Silva Rosa SC, Piotrowski S, Chapman D et al. A conserved MADS-box phosphorylation motif regulates differentiation and mitochondrial function in skeletal, cardiac, and smooth muscle cells. Cell Death Dis 2015; 6: e1944.

31 Kwong JQ, Lu X, Correll RN, Schwanekamp JA, Vagnozzi RJ, Sargent MA et al. The Mitochondrial Calcium Uniporter Selectively Matches Metabolic Output to Acute Contractile Stress in the Heart. Cell Rep 2015; 12: 15–22.

32 Luongo TS, Lambert JP, Yuan A, Zhang X, Gross P, Song J et al. The Mitochondrial Calcium Uniporter Matches Energetic Supply with Cardiac Workload during Stress and Modulates Permeability Transition. Cell Rep 2015; 12: 23–34.

33 Szabadkai G, Bianchi K, Varnai P, De Stefani D, Wieckowski MR, Cavagna D et al. Chaperone-mediated coupling of endoplasmic reticulum and mitochondrial Ca2+ channels. J Cell Biol 2006; 175: 901–911.

34 De Stefani D, Bononi A, Romagnoli A, Messina A, De Pinto V, Pinton P et al. VDAC1 selectively transfers apoptotic Ca2+ signals to mitochondria. Cell Death Differ 2011;: 1–7.

35 Zhao Y, Araki S, Wu J, Teramoto T, Chang YF, Nakano M et al. An Expanded Palette of Genetically Encoded Ca2+ Indicators. Sci N Y NY 2011; 333: 1888–1891.

36 Mughal W, Martens M, Field J, Chapman D, Huang J, Rattan S et al. Myocardin regulates mitochondrial calcium homeostasis and prevents permeability transition. Cell Death Differ 2018. doi:10.1038/s41418-018-0073-z.

37 Chen G, Ray R, Dubik D, Shi L, Cizeau J, Bleackley RC et al. The E1B 19K/Bcl-2-binding protein Nip3 is a dimeric mitochondrial protein that activates apoptosis. J Exp Med 1997; 186: 1975–1983.

38 Shimizu S, Konishi A, Kodama T, Tsujimoto Y. BH4 domain of antiapoptotic Bcl-2 family members closes voltage-dependent anion channel and inhibits apoptotic mitochondrial changes and cell death. Proc Natl Acad Sci U S A 2000; 97: 3100–3105.

39 Wilkins BJ, De Windt LJ, Bueno OF, Braz JC, Glascock BJ, Kimball TF et al. Targeted disruption of NFATc3, but not NFATc4, reveals an intrinsic defect in calcineurin-mediated cardiac hypertrophic growth. Mol Cell Biol 2002; 22: 7603–7613.

40 Wilkins BJ, Dai Y-S, Bueno OF, Parsons SA, Xu J, Plank DM et al. Calcineurin/NFAT coupling participates in pathological, but not physiological, cardiac hypertrophy. Circ Res 2004; 94: 110–118.

41 Wu X, Zhang T, Bossuyt J, Li X, McKinsey TA, Dedman JR et al. Local InsP3-dependent perinuclear Ca2+ signaling in cardiac myocyte excitation-transcription coupling. J Clin Invest 2006; 116: 675–682.

42 Chen Y, Decker KF, Zheng D, Matkovich SJ, Jia L, Dorn GW. A nucleus-targeted alternately spliced Nix/Bnip3L protein isoform modifies nuclear factor κB (NFκB)-mediated cardiac transcription. J Biol Chem 2013; 288: 15455–15465.

43 Chen Lf null, Fischle W, Verdin E, Greene WC. Duration of nuclear NF-kappaB action regulated by reversible acetylation. Science 2001; 293: 1653–1657.

44 Chen L-F, Williams SA, Mu Y, Nakano H, Duerr JM, Buckbinder L et al. NF-kappaB RelA phosphorylation regulates RelA acetylation. Mol Cell Biol 2005; 25: 7966–7975.

45 Kondo K, Klco J, Nakamura E, Lechpammer M, Kaelin WG. Inhibition of HIF is necessary for tumor suppression by the von Hippel-Lindau protein. Cancer Cell 2002; 1: 237–246.

46 Wu J, Liu L, Matsuda T, Zhao Y, Rebane A, Drobizhev M et al. Improved Orange and Red Ca 2+Indicators and Photophysical Considerations for Optogenetic Applications. ACS Chem Neurosci 2013; 4: 963–972.

47 Wu J, Prole DL, Shen Y, Lin Z, Gnanasekaran A, Liu Y et al. Red fluorescent genetically encoded Ca 2+indicators for use in mitochondria and endoplasmic reticulum. Biochem J 2014; 464: 13–22.

48 Wang NS, Unkila MT, Reineks EZ, Distelhorst CW. Transient expression of wild-type or mitochondrially targeted Bcl-2 induces apoptosis, whereas transient expression of endoplasmic reticulum-targeted Bcl-2 is protective against Bax-induced cell death. J Biol Chem 2001; 276: 44117–44128.

49 Perciavalle RM, Stewart DP, Koss B, Lynch J, Milasta S, Bathina M et al. Anti-apoptotic MCL-1 localizes to the mitochondrial matrix and couples mitochondrial fusion to respiration. Nat Cell Biol 2012; 14: 575–583.

50 Guan B, Wang T-L, Shih I-M. ARID1A, a factor that promotes formation of SWI/SNF-mediated chromatin remodeling, is a tumor suppressor in gynecologic cancers. Cancer Res 2011; 71: 6718–6727.

51 Perry RLS, Yang C, Soora N, Salma J, Marback M, Naghibi L et al. Direct interaction between myocyte enhancer factor 2 (MEF2) and protein phosphatase 1alpha represses MEF2-dependent gene expression. Mol Cell Biol 2009; 29: 3355–3366.

52 Du M, Perry RLS, Nowacki NB, Gordon JW, Salma J, Zhao J et al. Protein kinase A represses skeletal myogenesis by targeting myocyte enhancer factor 2D. Mol Cell Biol 2008; 28: 2952–2970.

53 Lee M, Lapham A, Brimmell M, Wilkinson H, Packham G. Inhibition of proteasomal degradation of Mcl-1 by cobalt chloride suppresses cobalt chloride-induced apoptosis in HCT116 colorectal cancer cells. Apoptosis Int J Program Cell Death 2008; 13: 972–982.

54 Alizadeh J, Zeki AA, Mirzaei N, Tewary S, Rezaei Moghadam A, Glogowska A et al. Mevalonate Cascade Inhibition by Simvastatin Induces the Intrinsic Apoptosis Pathway via Depletion of Isoprenoids in Tumor Cells. Sci Rep 2017; 7: 44841.

55 Pagiatakis C, Gordon JW, Ehyai S, Mcdermott JC. A Novel RhoA/ROCK-CPI-17-MEF2C Signaling Pathway Regulates Vascular Smooth Muscle Cell Gene Expression. J Biol Chem 2012; 287: 8361–8370.

56 Gordon JW, Pagiatakis C, Salma J, Du M, Andreucci JJ, Zhao J et al. Protein kinase A-regulated assembly of a MEF2middle dotHDAC4 repressor complex controls c-Jun expression in vascular smooth muscle cells. J Biol Chem 2009; 284: 19027–19042.

